# PDG-Arena: An ecophysiological model for characterizing tree-tree interactions in heterogeneous and mixed stands

**DOI:** 10.1101/2024.02.09.579667

**Authors:** Camille Rouet, Hendrik Davi, Arsène Druel, Bruno Fady, Xavier Morin

## Abstract

In the context of the ongoing climate and biodiversity crises, mixed forest stands are increasingly considered as a sustainable management alternative to monocultures. We developed a new individual-based and process-based forest growth model, PDG-Arena, to simulate mixed forest functioning and test ecophysiological interactions among trees in mixed stands. The model builds upon the validated ecophysiological stand-scale model CASTANEA and integrates tree competition for light and water. We evaluated the performance of PDG-Arena by comparing the simulated growth with annual dendrochronological growth data from 37 common beech and silver fir monospecific and mixed plots in the French Alps. PDG-Arena showed a slightly better performance than CASTANEA when simulating even-age and monospecific forests (r^2^ of 32.1 versus 29.5%). When using structure-diverse and species-diverse inventories, PDG-Arena performed better than CASTANEA in pure beech (38.3 versus 22.9%) and mixed stands (40.5 versus 36.3%), but not in pure fir stands (39.8 versus 42.0%). The new model also showed a significant positive effect of species mixing on gross primary production (+5.5%), canopy absorbance (+11.1%) and transpiration (+15.8%). Our results thus show that tree-level process-based models such as PDG-Arena, formally simulating interspecific interactions, can serve as a valuable tool to understand and simulate the carbon, light and water dynamics of mixed stands.

## 1. Introduction

Understanding how forest ecosystems function is a crucial step for developing forest management strategies adapted to the challenges of climate change (Bonan, 2008; Lindner et al., 2010; Trumbore et al., 2015) and more generally global change (González de Andrés, 2019). In this context, mixed forests, in comparison with monospecific stands, have received increasing attention due to their documented ability to maintain key ecosystem services while enhancing stand resilience (van der Plas et al., 2016; Seynave et al., 2018; Messier et al., 2022; del Río et al., 2022).

However, the ecophysiological functioning of mixed stands is still poorly understood (Forrester, 2014; For- rester and Bauhus, 2016). In particular, even though species mixing seems on average to increase stand productivity in comparison to monospecific stands (a phenomenon known as overyielding) (Liang et al., 2016; Zhang et al., 2012; Vilà et al., 2007; Forrester and Bauhus, 2016; Piotto, 2008), this trend depends on stand structure and species composition (Zhang et al., 2012; Ratcliffe et al., 2015), as well as abiotic conditions (Ratcliffe et al., 2016; Toïgo et al., 2015). Regarding the effect of tree species richness on the resistance of stands to drought episodes, the literature shows heterogeneous results (Grossiord, 2018). Indeed, the direction of the effect seems to depend on species composition - and particularly on the species respective strategies in reaction to soil water deficit (Pretzsch et al., 2013; Mas et al., 2024; Jourdan et al., 2020) - as well as on environmental conditions (Grossiord et al., 2014; Forrester et al., 2016; Pardos et al., 2021).

Stand structure, particularly tree density and size variability, can act as a confounding factor in the diversity-functioning relationship (Metz et al., 2016; Dă nescu et al., 2016; Cordonnier et al., 2019; Zeller and Pretzsch, 2019). To better understand the processes underlying these relationships, it is therefore important to separate the effects of mixing related to differences in stand structure (age, size, diameter) from those related to differences in the physiological functioning of species (crown architecture, water strategy, nutrient use, etc.; see Forrester and Bauhus, 2016).

Furthermore, the interactions observed in a mixture may be of various kinds (Forrester et al., 2016), which could give rise to contradictory effects. For example, an increase in the amount of light captured in mixtures - e.g., through crown complementarity and plasticity, see Jucker et al. (2015) - could lead to an increase in gross primary production, but also in transpiration, with a potentially negative effect on available soil water (Jucker et al., 2014). Forrester (2014) proposed a conceptual model to account for the mechanisms of interaction between diversity, functioning and environment. In this framework, interspecific interactions resulting in reduced competition for a given type of resource generate beneficial effects for individuals when this resource becomes scarce.

Assessing and predicting the functioning of mixed stands therefore requires detailed knowledge of interspecific interactions. This knowledge must be based on interactions between individuals and on the ecophysiological processes underlying these interactions, i.e., the processes determining competition for light, water and nutrients (Pretzsch et al., 2017; Grossiord, 2018). This knowledge is all the more necessary as abiotic and biotic conditions are and will be transformed by global change (Ammer, 2019).

Although experimental and observational systems are necessary for studying the diversity-functioning relationship in forests, they are limited by their sample size, measurement completeness and number of confounding factors that can be controlled (Bauhus et al., 2017). Modeling can virtually overcome these limitations, subject to the assumptions contained in the model, which depend to a large extent on our ecological knowledge as well as on the availability of climatic, pedological, silvicultural and physiological data. The modeling approach has been used to put forward hypotheses to explain overyielding in mixing. For example Morin et al. (2011) showed with simulations that overyielding could be explained by the diversity of species traits related to shade-tolerance, maximum height and growth rate (although other explanations could not be ruled out). Simulations also make it possible to virtually assess the stability of the productivity of forest mixtures while testing numerous community compositions (Morin et al., 2014), even under unprecedented climatic conditions (Jourdan et al., 2021).

The literature (Korzukhin et al., 1996; Cuddington et al., 2013; Morin et al., 2021) depicts a spectrum ranging from empirical models, which are based on relationships calibrated from observations between final variables such as productivity and explanatory variables (e.g., rainfall, sunshine), to process-based models where final variables are computed using explicit elementary processes (e.g., photosynthesis, transpiration, phenology). For some authors (Fontes et al., 2010; Cuddington et al., 2013; Korzukhin et al., 1996), process-based models seem more relevant for simulating ecosystem functioning undergoing climate change because they can theoretically be applied to a larger range of environmental conditions than empirical ones. As a result, they now play an important role in research on the ecophysiological functioning and prediction of forest dynamics (Gonçalves et al., 2021; Barbosa et al., 2023). However, compared to empirical models, process-based models are more difficult to parameterize and rely on more assumptions about the ecological functioning of forests (e.g., the hypothesis that growth is primarily driven by photosynthetic activity, Fatichi et al., 2014). When it comes to simulating mixed stands, models that simulate elementary processes are expected to reproduce the mechanisms that lead to interspecific interactions, bringing us closer to understanding them (Forrester and Bauhus, 2016).

Among process-based models, a distinction is made between individual-based models, e.g., Jonard et al. (2020), and stand-scale models, e.g., Dufrêne et al. (2005). Several diversity-functioning studies in forests have highlighted the importance of tree-tree interactions in defining the nature of interspecific interactions at stand level (Trogisch et al., 2021; Jourdan et al., 2020; Guillemot et al., 2020; Jucker et al., 2015). Thus, the individual scale appears relevant for representing the key mechanisms that govern the functioning of mixed forests (Porté and Bartelink, 2002). Finally, process-based and individual-based models have the ability to distinguish the effects of competition between individuals of different species (mixing effect) and the effects of competition between individuals of different sizes (structure effect). So far, few models are able to simulate mixed stands by taking advantage of both physiological mechanisms and the individual scale (Reyer, 2015; Pretzsch et al., 2015).

Here we present a new individual-based and process-based forest growth model, PDG-Arena (the arena represents the stand, a place where trees compete and more generally interact). Our model was developed to observe the stand scale properties that emerge when trees of different species and size compete in a given environment. It was therefore built: (i) from elementary physiological processes using the stand-scale model CASTANEA (Dufrêne et al., 2005) and (ii) by integrating interactions among trees, notably competition for light and water.

The performance of PDG-Arena was evaluated using annual growth data from a monitoring network of monospecific and multispecific stands of common beech (*Fagus sylvatica* L.) and silver fir (*Abies alba* Mill.). Firstly, we tested whether PDG-Arena, despite increased complexity, accurately reproduces the performance of CASTANEA when both models are run under comparable conditions. Secondly, we evaluated PDG-Arena’s performance in different conditions in terms of stand structure and species diversity. Lastly, using PDG-Arena, we evaluated the effect of species mixing on carbon, light and water processes.

## 2. Materials & Methods

### 2.1. Model description

#### 2.1.1. From CASTANEA to PDG-Arena

PDG-Arena was designed as an extension of PDG (which stands for Physio-Demo-Genetics, Oddou-Muratorio and Davi, 2014), an individual-based and spatially explicit model that combines: (1) the process-based model CASTANEA to simulate tree ecophysiology, (2) demographic processes allowing to model tree survival and reproduction and (3) a quantitative genetics simulation module accounting for the heritability and intraspecific diversity of key life history trait of the CASTANEA model. While PDG is built with the idea of simulating the evolutionary dynamics of functional traits of importance for adaptive forestry in regular monospecific stands (Lefèvre et al., 2014), PDG-Arena is designed to simulate ecological interactions between trees in diverse, multispecific stands.

CASTANEA is an ecophysiological forest growth model that simulates the dynamics of homogeneous stands (Figure 1a). Among others, it has been parameterized and validated on common beech (Fagus sylvatica L., Dufrêne et al., 2005) and silver fir (Abies alba Mill., Davi and Cailleret, 2017). CASTANEA is composed of five equal-sized leaf layers that perform photosynthesis based on stomatal conductance and on the level of radiation received by each layer, which is determined using a horizontally homogeneous multi-layer radiation model. The resulting gross primary production, minus autotrophic respiration, is then allocated into the leaf, fine root, coarse root, branch, trunk and reserves compartments (Davi et al., 2009). The amount of leaf transpiration is determined by net radiation, stomatal conductance as well as ambient temperature and vapor pressure deficit. The stomatal conductance, limiting photosynthesis and transpiration, is controlled by soil water deficit (using the critical threshold of relative extractable water *REW*_*c*_ in Granier et al., 1999). Lastly, leaf surface growth is controlled by day length and mean temperature. The temporal scale of the processes in CASTANEA is the same as that of PDG-Arena, as shown in Table 1.

**Table 1:** Temporal and spatial scales of physical and physiological processes in PDG-Arena.

|  | Tree level | Stand level |
| --- | --- | --- |
| **Hourly level** | Photosynthesis<br>Respiration<br>Crown transpiration<br>Crown evaporation | Ray casting<br>Soil evaporation |
| **Daily level** | Water interception<br>Leaf phenology<br>Carbon allocation | Water balance |
| **Yearly level** | Tree growth |  |

**Figure 1:**
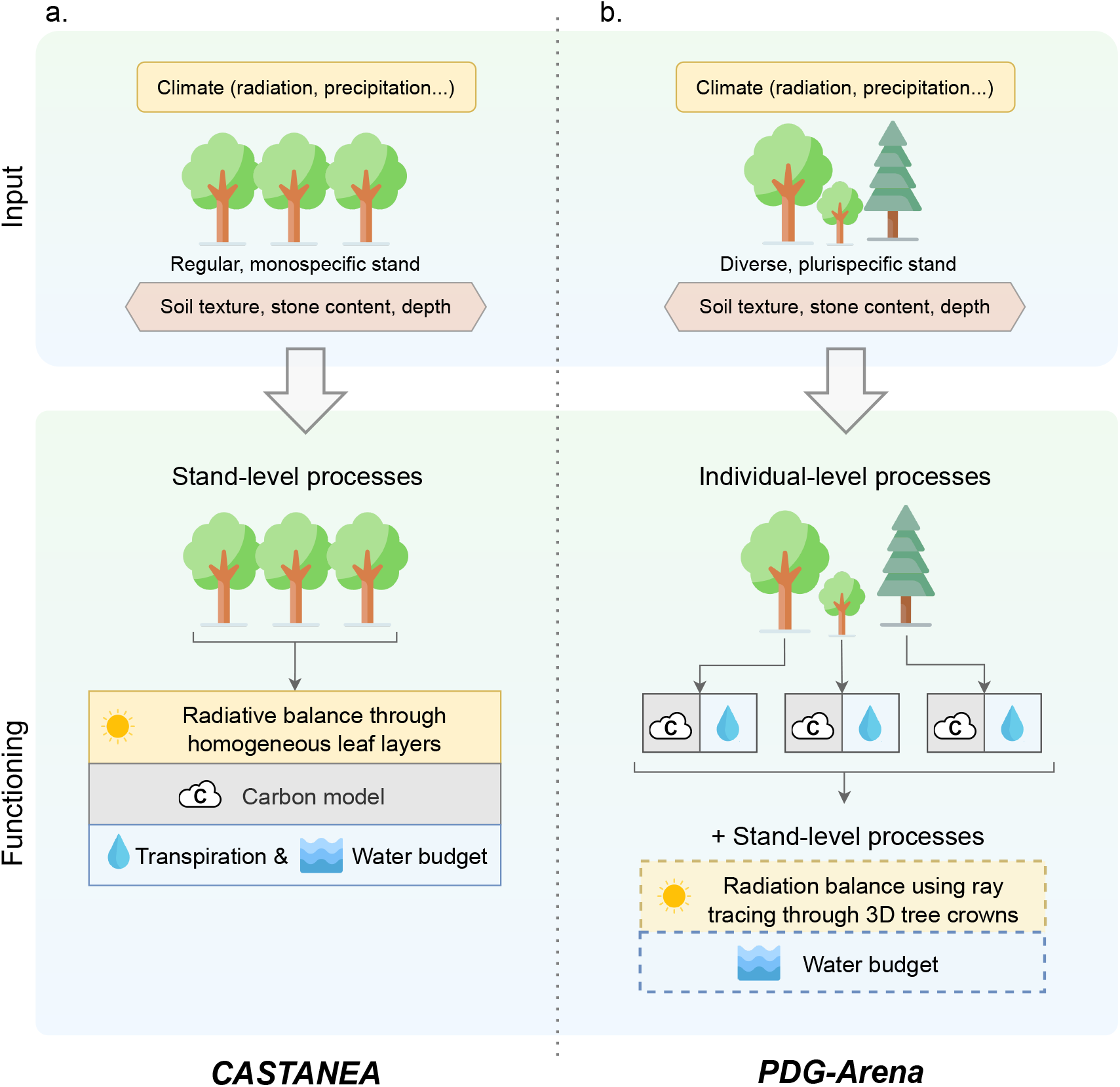
Conceptual diagram of the (a) CASTANEA and (b) PDG-Arena forest growth models input and functioning. CASTANEA and PDG-Arena respectively simulate the growth of regular monospecific stands and (potentially) diverse multispecific stands. In CASTANEA, all processes, including radiation balance, carbon fluxes, transpiration of trees and soil water budget occur at the stand level, on horizontally homogeneous leaf layers. PDG-Arena takes advantage of CASTANEA carbon and transpiration processes but performs them at the tree level, while a water budget is computed at the stand level. Its radiative balance is handled by the SamsaraLight library which casts light rays through a 3D representation of tree crowns. Processes involving competition between trees in PDG-Arena are shown in dashed boxes.

The existing model PDG considers isolated abstract trees, simulating the dynamics of each of them using stand-scale processes of CASTANEA. All quantitative physiological variables in CASTANEA and in PDG are expressed on a per area basis: e.g., the gross primary production is expressed in gC/m^2^. The first improvement of PDG-Arena over PDG is that the physiological processes simulate tree functioning instead of stand functioning (Figure 1b). To do so, physiological processes are related to the projected area of the individual crowns rather than to the stand area. This paradigm shift implied changing the definition of some variables. As depicted in Figure 2, the Leaf Area Index (LAI) is now defined for each tree as the amount of leaf surface of a tree per m^2^ of soil under its crown. While the stand LAI in CASTANEA depends on gap fraction, individual tree LAI in PDG-Arena does not: the LAI of a tree only accounts for its leaf surface and its crown projection surface. The same reasoning applies to other physiological variables, such as carbon uptake, water transpiration, absorbed radiation, etc. Also, the Leaf Mass Area (LMA), as it depends on the amount of light intercepted by neighboring trees, is computed at the individual level in PDG-Arena according to the vertical profile of the leaf area of neighboring trees (see Appendix B.1).

**Figure 2:**
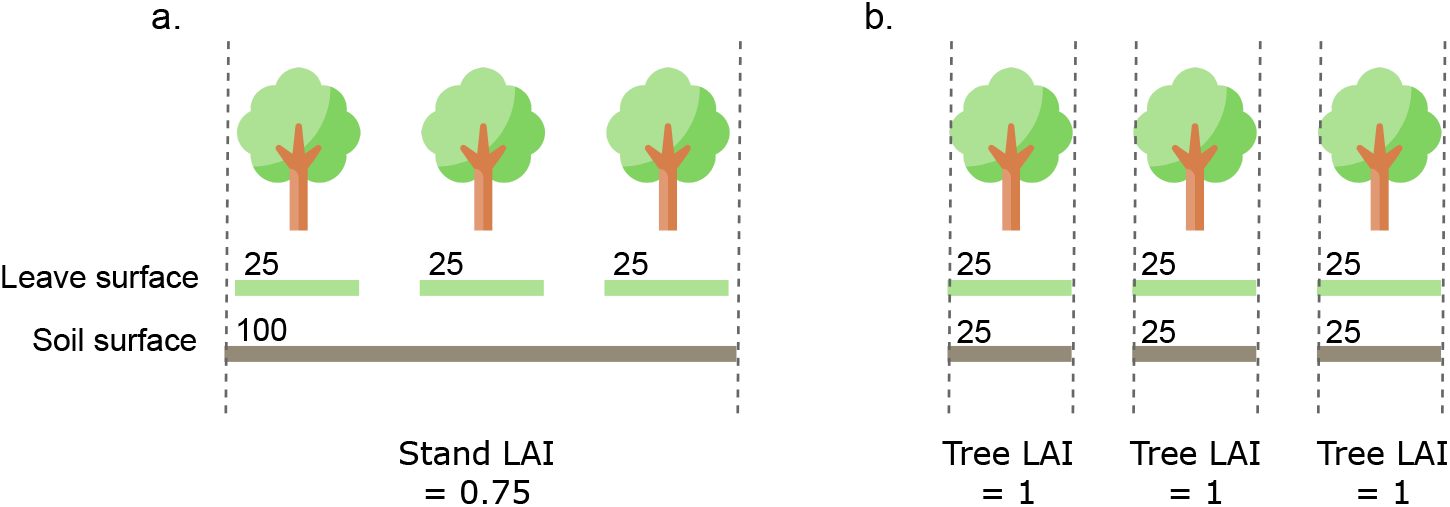
Difference in the representation of Leaf Area Index (LAI) between (a.) the stand-scale model CASTANEA and (b.) the individual-based model PDG-Arena. Values of leaf surface, soil surface and LAI are arbitrary.

The second improvement of PDG-Arena over PDG is that it integrates mechanisms of competition for light and water between neighboring trees (see Figure 1b) by: (i) making trees share the same stand soil water pool and (ii) simulating irradiance at tree level using a ray tracing model.

#### 2.1.2. Competition for water

Competition for water is a crucial element in the dynamics of mixed stands. We modeled competition for water symmetrically between individuals, i.e., trees in the same plot all draw from the same water reservoir without spatial differentiation, either horizontal (distance between individuals) or vertical (depth).

Every day of simulation, the stand-level volume of precipitation is divided into a fraction that does not interact with the canopy – i.e., that falls directly to the ground – and another fraction that reaches the canopy. The fraction that interacts with the canopy is given by the proportion of soil that is directly under any tree crown. Then, this fraction of precipitation is distributed among trees according to their respective leaf surface. For each tree, a calculation of drip, runoff, and precipitation passing through the crown is performed using the same equation as in CASTANEA (Dufrêne et al., 2005). Transpiration and crown evaporation of trees are calculated individually at hourly time steps using the Penman-Monteith equation (Monteith, 1965), taking into account the energy absorbed by individual crowns (see section 2.1.3). Stand soil evaporation is computed hourly and homogeneously along the plot, following equations of CASTANEA (Dufrêne et al., 2005). Evapotranspiration from understorey vegetation is ommited.

Considering drip, runoff and water passing through the crowns on the one hand, and tree transpiration, canopy and soil evaporation and drainage on the other, a water balance is computed each day at the stand level (Table 1 and Figure 1b). Therefore, soil water status (soil moisture, litter moisture and soil potential) is the same for every tree within a plot on any given day.

#### 2.1.3. Competition for light

Competition for light in PDG-Arena is performed using SamsaraLight, a ray tracing library derived from Courbaud et al. (2003) and maintained on the Capsis modeling platform. The integration of SamsaraLight with the physiological model CASTANEA (which is partly inspired from the approach in the HETERO-FOR model, Jonard et al., 2020) is described here. Light conditions are evaluated both in the PAR (photosynthetically active radiation) domain and in the NIR (near infrared radiation) domain. For each domain, SamsaraLight generates a set of diffuse and direct beams, and computes their interception by tree crowns and soil cells. The simulated energy absorbed by crowns is then temporally distributed at the hourly scale. The energy absorbed by a crown is distributed among its five leaf layers, which are part of the CASTANEA model for each tree.

##### Definition of crowns

Each tree is represented by a crown occupying a volume in space and is defined by the following variables:

- the height of the tree *h*;
- its crown base height, *hcb*;
- its crown radius *crownRadius*;
- its shape, which is considered conical in the case of silver fir and ellipsoidal in the case of common beech (shapes are vertically bounded by *h* and *hcb* and horizontally bounded by *crownRadius*);
- its leaf area density at period of full vegetation, denoted *LAD*, in m^2^ of leaf per m^3^ of crown volume;
- its attenuation coefficient *k*;
- its clumping index Ω defining the aggregation of the leaves inside the crown.

Tree *h* and *hcb* values are model inputs (see section 2.2). Tree crown radius is estimated using an allometric relationship based on species and diameter at breast height (DBH):

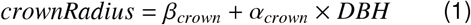

*α*_*crown*_ and *β*_*crown*_ are species dependent parameters estimated on site at Mont Ventoux (unpublished data from one of the authors, H. Davi). Ω is species dependent and was measured on Mont Ventoux sites by Davi et al. (2008b). The attenuation coefficient *k* depends on species, radiation domain, type of radiation (direct, diffuse) and beam height angle. Its value is determined using reverse-engineering of SAIL (the radiation sub-model in CASTANEA) as described in Appendix B.2.

The *LAD* of a tree is the ratio of its leaf area to its crown volume. The leaf area of a given tree *i* (denoted *LA*_*i*_) is determined using the stand leaf area at full vegetation (*LA*_*stand*_, which is a simulation input, see section 2.2). For every tree, its fraction of leaf area over stand leaf area is proportional to its theoretical leaf area *LA*_*th*_:

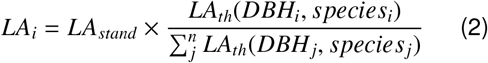

*LA*_*th*_ is given by an allometric equation based on DBH and species from Forrester et al. (2017b):

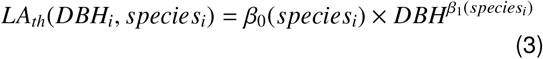

During the radiation balance computation, each tree LAD is at its maximum. However, a fraction of the absorbed radiations per tree is removed daily depending on their current phenological state (see Appendix B.4).

##### Ray casting

SamsaraLight generates two sets of beams. Firstly, diffuse rays are generated in all directions, using a 5°discretization. Secondly, direct rays are generated to follow the hourly trajectory of the sun for one virtual day per month. Each set of beams contains the energy of the entire year for both diffuse and direct radiations. The stand plot is subdivided into square cells of 1.5 m width. All beams are replicated for each ground cell, aiming at the center of the cell.

Once all the rays have been created, SamsaraLight performs the ray casting as described in Courbaud et al. (2003). For each ray, its energy is attenuated when it crosses a crown. The proportion of energy transmitted follows the formulation of the Beer-Lambert law:

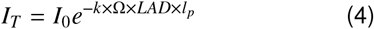

where *l*_*p*_ is the path length of the ray in the crown and *I*_0_ is the energy of the beam before it intercepts the crown. Then, the energy absorbed by a crown *I*_*A*_ is the complement of the transmitted energy:

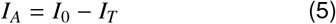

Note that SamsaraLight does not take directly into account the reflection of light - which causes a loss of energy in the sky and a reabsorption of the energy reflected on the ground. These phenomena are taken into account when calculating the attenuation coefficient.

After interception by a crown, the ray continues its course until it reaches either a new crown or a ground cell to which the remaining energy is transmitted. A proportion of absorbed radiation *ϵ* is uniformly removed from soil cells to represent the light extinction from trunks, assuming a random arrangement of trees:

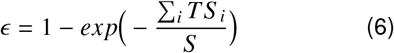

where *S* is the stand area and _*i*_ *TS* _*i*_ is the sum of the trunk shade surface of individual trees. *TS* _*i*_ depends on the DBH and height of each tree *i* (supposing a cylindrical shape of the trunk), as well as on the hourly sun angle *β*(*h*):

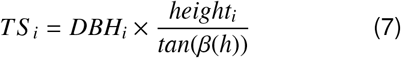

At the end of the ray casting, we know for each crown and each soil cell the amount of direct and diffuse energy received over a year.

##### Computation of hourly absorbed energy

The hourly absorbed radiation of any element is then computed using the ray casting on the one hand and the hourly incident radiation on the other hand.

For each absorbing element *i* (a soil cell or a tree crown) and for each type of radiation (direct/diffuse, PAR/NIR), the energy it absorbs at hourly scale is given by the hourly incident radiation *gr*(*h*) and the fraction of energy absorbed annually by this element, *I*_*Ay*_(*i*), divided by the total energy absorbed by all elements *j* over the year:

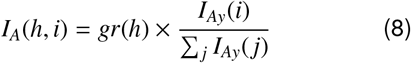

The value of *I*_*A*_(*h, i*) has then to be amended because the ray casting uses values of LAD that assume trees are at their period of full vegetation. A surplus of energy is then removed afterward from each tree according to their daily level of leaf development. This surplus is redistributed into other trees and soil cells, as described in Appendix B.4.

##### Distribution among layers

Within a real-life tree, some leaves can receive a large amount of light - which leads to a saturation of the photosynthesis capacities - while others are in the shade. The saturation phenomenon (and more generally the concavity of the absorbed light-photosynthesis relation) forbids calculating photosynthesis by considering an average level of light absorption for the whole canopy: this would bias upwards the estimation of photosynthesis (Leuning et al., 1995). In CASTANEA, the energy absorbed by the canopy is therefore distributed into five layers of leaves, in which the absorbed energy is assumed to be relatively homogeneous. The layers are themselves divided between leaves under direct light (called sun leaves) and leaves in the shade. The distribution of energy into the different layers is described in Appendix B.3.

### 2.2. Data set

To evaluate the simulations, we used an existing data set (GMAP forest plot design, Jourdan et al., 2019, 2020) composed of 39 beech, fir and beech-fir plots sampled between 2014 and 2016. Plots are distributed on three sites from the French pre-Alps (Bauges, Ventoux, Vercors), which are described in Table 2. They consist in a 10 m radius area in which the position, height, crown base height, age, diameter and species of each tree with a DBH greater than 7.5 cm were collected once.

**Table 2:** Characteristics of the stands used to evaluate the model. Mean value and standard deviation for each site (Bauges, Ventoux, Vercors) and composition (Mixed, Beech, Fir) are shown for variables: number of stands, altitude (in m), mean diameter at breast height per stand (in cm), density (in stem/ha), basal area (in m^2^/ha), proportion of beech basal area (in %), mean age per stand, Leaf Area Index (no unit).

| Site / Composition | N | altitude | mean DBH | density | basal area | % beech | mean age | LAI |
| --- | --- | --- | --- | --- | --- | --- | --- | --- |
| Bauges | 10 | 1100 ± 101 | 28.7 ± 6.7 | 1030 ± 685 | 72 ± 14 | 53 ± 43 | 89 ± 16 | 5.6 ± 0.2 |
| Vercors | 14 | 1250 ± 101 | 32.3 ± 8.6 | 657 ± 275 | 56 ± 14 | 53 ± 38 | 118 ± 40 | 5.6 ± 0.3 |
| Ventoux | 15 | 1250 ± 126 | 22.1 ± 6.3 | 1450 ± 623 | 57 ± 13 | 50 ± 40 | 105 ± 47 | 3.2 ± 0.3 |
| Mixed | 13 | 1200 ± 131 | 26.2 ± 7.3 | 1080 ± 465 | 64 ± 13 | 46 ± 10 | 101 ± 29 | 4.7 ± 0.5 |
| Beech | 14 | 1230 ± 118 | 26.7 ± 10.3 | 1200 ± 794 | 56 ± 14 | 97 ± 5 | 119 ± 35 | 4.7 ± 1.2 |
| Fir | 12 | 1190 ± 139 | 29.8 ± 7.4 | 867 ± 578 | 62 ± 18 | 5 ± 7 | 94 ± 50 | 4.7 ± 1.3 |
| All | 39 | 1210 ± 126 | 27.5 ± 8.4 | 850 ± 632 | 60 ± 15 | 51 ± 39 | 105 ± 39 | 2.9 ± 1.2 |

Out of 1177 stems, 731 were cored to assess the growth dynamics over the 18-year period 1996-2013 (Jourdan et al., 2019). Growth of non-cored stems was inferred on the assumption that basal area increment over basal area was constant for a given species and site. To be comparable with the model output, basal area increments were converted into wood volume increments. To do that, we inferred past tree heights by using values of past DBH and the relationship between measured height and DBH. Past DBH were reconstructed using basal area increments and measured DBH. Then, a model was fitted on trees of the same species and site to evaluate the relationship between measured height and DBH (see Appendix A). This model was used to compute past height based on reconstructed past DBH.

Wood volume increments were computed by multiplying each tree basal area increment with its inferred past height and Φ, a form factor coefficients which takes into account the non-cylindrical shape of the trunks (Deleuze et al., 2014). On the one hand, PDG-Arena was evaluated using wood volume increments at individual scale. On the other hand, we used the wood volume increments at stand scale to evaluate both PDG-Arena and CASTANEA.

Hourly climate data (temperature, global radiation, wind speed, precipitation and relative humidity) were obtained from the 8 km scale SAFRAN reanalysis data set (Vidal et al., 2010) for the three sites and temperatures were adapted to each stand altitude using an adjustment of 0.6 °C/100m (Rolland, 2003). Soil texture, depth and stone content were obtained for every stand (data from one of the authors, X. Morin, see section 2.5 for accessibility).

The LAI of the stands were retrieved using each plot coordinates and the 1 km resolution SPOT/PROBA-V remote sensing data set (Baret et al., 2013). We computed the average value of the yearly maximum LAI observed over the 1999-2013 period (see section 2.5 for accessibility)

### 2.3. Simulation plan

Using field inventories, we generated three sets of virtual inventories for PDG-Arena, following three levels of abstraction, denoted RM, R and O. The first set represents regularized monospecific inventories (RM): for each species of each stand, we generated a new inventory with equally spaced trees of the same species, age, diameter and height. For mixed stands, the simulation results using RM inventories were assembled relatively to the proportion of each species basal area. RM inventories can then be used to simulate the growth of multispecific stands while ignoring species interactions. The second set represents regularized inventories (R), in which trees of different species can coexist but trees of the same species share the same age, diameter and height. Trees in R inventories are regularly spaced in a random order, independently of the species. Lastly, original inventories (O) include the information of the real life data set, that is: species, position, diameter and height of every individual trees. For each type of inventories representing the same stand (regularized or not, with or without species interactions), the mean quadratic diameter, volume per tree and tree age per species and the basal area were conserved.

CASTANEA was used as a reference model to evaluate the performance enhancement brought by PDG-Arena. We used RM inventories for CASTANEA’s stand-scale simulations. It is to be noted that, contrary to PDG-Arena, CASTANEA does not account for the stand slope. Therefore, when comparing CASTANEA and PDG-Arena results (section 3.1), the slope was put to zero in PDG-Arena inventories. In the other situations (sections 3.2 and 3.3), the slopes of the inventories simulated using PDG-Arena were those of the field data.

To sum up, we simulated the growth of 39 stands over the 18-year period 1996-2013, considering four situations: RM, R and O inventories with PDG-Arena and RM inventories with CASTANEA. Tree reproduction and intraspecific diversity, which are characteristics of PDG and therefore PDG-Arena, were switched off for these simulations.

### 2.4. Model evaluation

To evaluate the similarity between each modeling situation, we used the gross primary production (GPP) as CASTANEA and PDG-Arena are carbon-based models. We computed the coefficient of correlation (r, from −1 to 1) for the simulated GPP per stand between the four situations.

To evaluate the performance of the models against field measurements, we used the simulated wood volume increment per stand. We computed the Mean Absolute Percentage Error (MAPE) and the coefficient of determination (r^2^, from 0 to 1) between simulations and measurements. A low MAPE indicates that simulated wood production is on average close to measured production. An r^2^ close to 1 shows a good capacity of the model to predict stand production variability. Additionally, PDG-Arena with O inventories was evaluated at the individual scale, by computing the r^2^ and MAPE of the simulated versus measured wood volume increment per tree for each group of the same site, type of stand (beech, fir of mixed) and species.

Lastly, we computed the net mixing effect (NME) to assess the extent of the simulated physiological processes that can solely be attributed to species mixing. Following the computation of the net biodiversity effect by Loreau (2010), we defined the NME as the difference for a variable between its observed value in mixed stands and its predicted value based on the hypothesis that there is no complementarity effect between species. Here, we compared the value of a simulated variable with PDG-Arena using the R and RM inventories (i.e., with and without species interactions). NME was evaluated on GPP, canopy absorbance, transpiration rate and maximum water shortage (defined as the maximum difference reached during simulation between the current and full useful reserve, in mm). We chose the maximum water shortage because, in comparison to the relative extractable water (REW) because it is expressed in absolute and is therefore independent of the site depth. NME was tested against the null hypothesis using a two-sided Wilcoxon signed rank test.

### 2.5. Access to model and data

The PDG-Arena model is part of the Capsis framework, which aims to facilitate collaborative and shared software development for forest science (Dufour-Kowalski et al., 2012). PDG-Arena is an extension of the Physio-Demo-Genetics model, whose code is only accessible to members of the Capsis community for testing purposes. PDG-Arena can be run through Capsis either by script or via a graphical interface, using a formalized inventory (describing each tree of the plot, the plot and soil characteristics and physiological, demographic and genetic options), as well as a CASTANEA-adapted climate file.

A repository is made accessible on the Zenodo platform (Rouet et al., 2024), containing:

- the script used for the generation of inventories;
- the inventories used is the simulations;
- two data sets describing the LAI and soil textures used in the simulations;
- the results data set (including formatted data of simulations and measurements);
- the script used for the analysis of the result data set;

Raw simulation results are accessible on demand.

## 3. Results

### 3.1. Comparison of PDG-Arena and CASTANEA

Using regular and monospecific inventories (RM), CASTANEA and PDG-Arena showed similar predictions for the stand-level GPP, with a coefficient of correlation at 99.8%. However, the GPP simulated by PDG-Arena was in average 4.2% greater than that of CASTANEA (Figure 3). As shown in Table 3, which compares the 4 modeling situations based on the coefficient of correlation, simulations from PDG-Arena was closer to those of CASTANEA when using regularized inventories (R) on the one hand and when using regularized monospecific inventories (RM) on the other hand.

**Table 3:** Matrix of similarity between simulated GPP from CASTANEA and PDG-Arena using different types of inventories: ‘RM’ (regularized and monospecific, i.e., without species interactions), ‘R’ (regularized, but with species interactions) and ‘O’ (original inventories). Similarity is expressed using the correlation coefficient (in %) of the simulated gross primary production for the 39 stands over the 1996-2013 period.

|  | CASTANEA<br>(RM) | PDG-Arena<br>(RM) | PDG-Arena<br>(R) | PDG-Arena<br>(O) |
| --- | --- | --- | --- | --- |
| CASTANEA (RM) | 100.0 | - | - | - |
| PDG-Arena (RM) | 99.8 | 100.0 | - | - |
| PDG-Arena (R) | 99.3 | 99.5 | 100.0 | - |
| PDG-Arena (O) | 97.7 | 98.5 | 99.0 | 100.0 |

**Figure 3:**
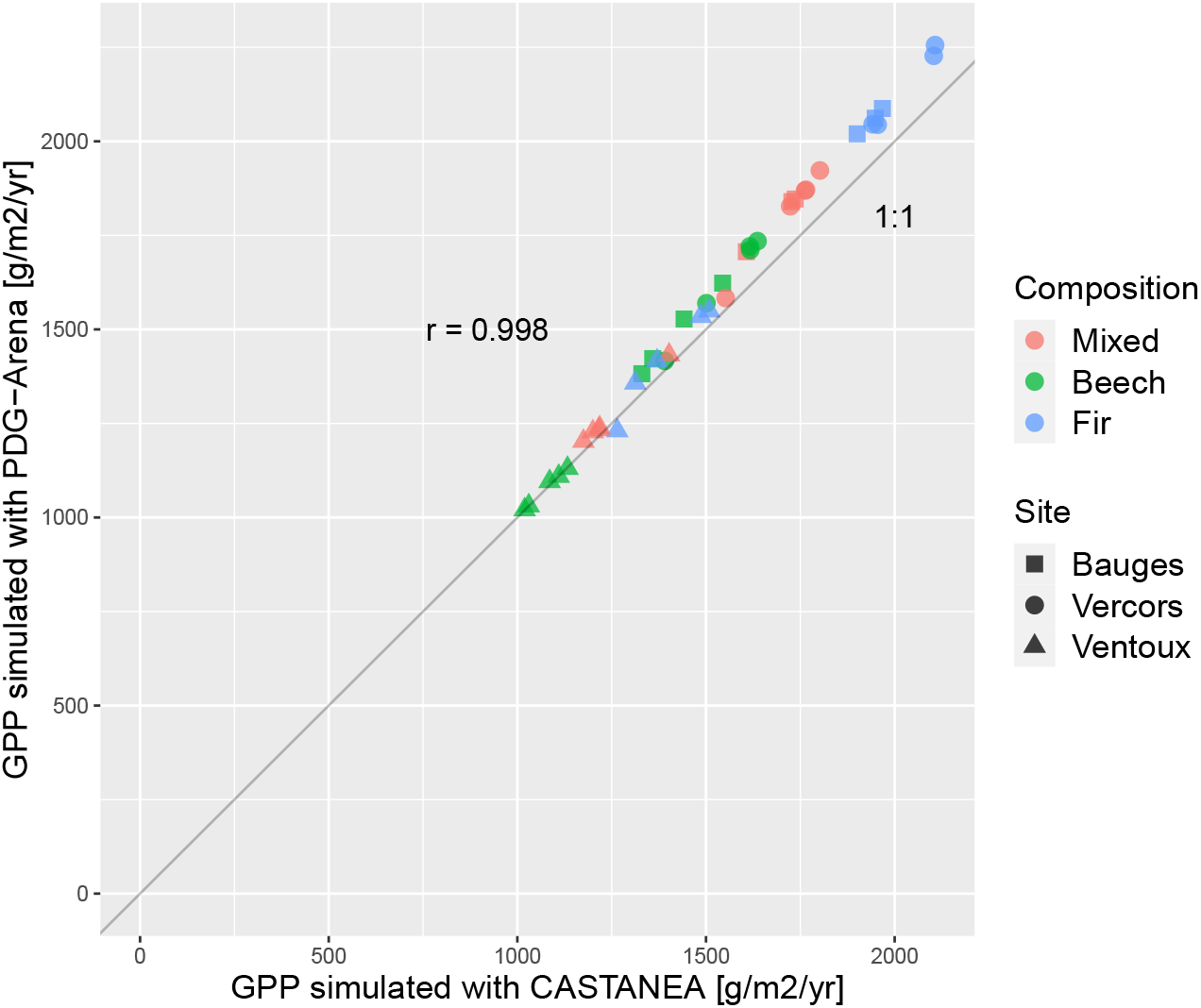
Gross primary production (GPP) per stand simulated by PDG-Arena and CASTANEA. Regularized monospecific inventories (RM) were used. *r* is the correlation coefficient.

### 3.2. Model performance

The simulated versus measured stand wood volume increment for the 39 stands are reported for the CASTANEA model using RM inventories and for the PDG-Arena model using O inventories in Figure C.6. Two fir stands from the Bauges site, denoted haut_sp_2 and bas_sp_4, stand out from the point cloud, with measured growths of 1995 and 1562 cm^3^/m^2^/year, while the simulated growth did not exceed 973 cm^3^/m^2^/year for CASTANEA and PDG-Arena. In addition, simulations using values of LAI measured in 2022 using Terrestrial Laser Scanning (unpublished data from one of the author, C. Rouet) were done and showed the same discrepancy with growth measurements for these two stands, indicating that the LAI measurement is not the problem. As the inclusion of these two stands in the analysis affects the overall results, they were discarded from the following analysis (see Table C.6 for the performance analysis that includes all stands).

Simulation performances of CASTANEA and PDG-Arena against measured wood volume increments per stand are reported in Table 4. The MAPE was close between models and types of inventories, ranging from 30.1 to 33.1% in mixed stands, 53.9 to 57.9% in beech stands and 29.6 to 33.7% in fir stands. Considering the 37 stands, performances were close between CASTANEA and PDG-Arena on comparable inventories, i.e., RM inventories, with a slight advantage for PDG-Arena (r^2^ 32.1% vs 29.5%). Using O inventories, PDG-Arena performed better than CASTANEA on RM inventories (r^2^ 34.2 vs 29.5%).

**Table 4:** Evaluation of the performances of PDG-Arena and CASTANEA on the 37 stands. Coefficient of determination (r^2^, in %) and Mean Absolute Percentage Error (MAPE, in %) were computed for the simulated versus measured yearly wood volume increment per stand over the period 1996-2013. Inventories are characterized as: ‘RM’ (regularized and monospecific, i.e., without species interactions), ‘R’ (regularized, but with species interactions) and ‘O’ (original inventories).

| Set | Model | Inventories | $r^2$ | MAPE |
| --- | --- | --- | --- | --- |
| All stands | CASTANEA | RM | 29.5 | 40.6 |
|  | PDG-Arena | RM | 32.1 | 40.5 |
|  | PDG-Arena | R | 32.5 | 41.8 |
|  | PDG-Arena | O | 34.2 | 40.4 |
| Mixed | CASTANEA | RM | 36.3 | 30.1 |
|  | PDG-Arena | RM | 37.6 | 30.7 |
|  | PDG-Arena | R | 36.3 | 33.1 |
|  | PDG-Arena | O | 40.5 | 31.5 |
| Beech pure | CASTANEA | RM | 22.9 | 55.3 |
|  | PDG-Arena | RM | 25.0 | 57.4 |
|  | PDG-Arena | R | 24.7 | 57.9 |
|  | PDG-Arena | O | 38.3 | 53.9 |
| Fir pure | CASTANEA | RM | 42.0 | 33.7 |
|  | PDG-Arena | RM | 51.9 | 29.6 |
|  | PDG-Arena | R | 50.1 | 30.4 |
|  | PDG-Arena | O | 39.8 | 33.0 |

Activation of species interactions in PDG-Arena (R vs RM inventories) slightly decreased the performance for mixed stands (r^2^ 36.3% vs 37.6%, MAPE 33.1% vs 30.7%). Using original instead of regularized inventories (O vs R), PDG-Arena displayed an improved performance on mixed (r^2^ 40.5 vs 36.3%, MAPE 31.5 vs 33.1%) and beech (r^2^ 38.3 vs 24.7%, MAPE 53.9 vs 57.9%) stands but a lower performance on fir stands (r^2^ 39.8 vs 50.1%, MAPE 39.8 vs 33.0%).

Figure C.7 shows the simulated versus measured wood volume increment at the tree scale using PDG-Arena and original inventories (O). The r^2^ ranged from 20 to 64% depending on the set of trees, with a mean at 47%. The MAPE ranged from 50 to 146%, with a mean of 82% (Table C.7).

### 3.3. Mixing and structure effects

GPP and canopy absorbance simulated by PDG-Arena in mixed stands are represented in Figure 4 for RM, R and O inventories. Additionally, Figure C.8 shows the yearly transpiration rate and maximum water shortage. Comparison of simulations with R and RM inventories showed a positive net mixing effect of 5.5% on GPP (1665 vs 1578 gC/m2/year; p-value < 0.001), of 11.1% on canopy absorbance (0.452 vs 0.407; p-value < 0.001), of 15.8% on canopy transpiration (234 vs 202 mm/year; p-value < 0.001) and of 13.7% on maximum water shortage (92.5 vs 81.3 mm; p-value < 0.001).

**Figure 4:**
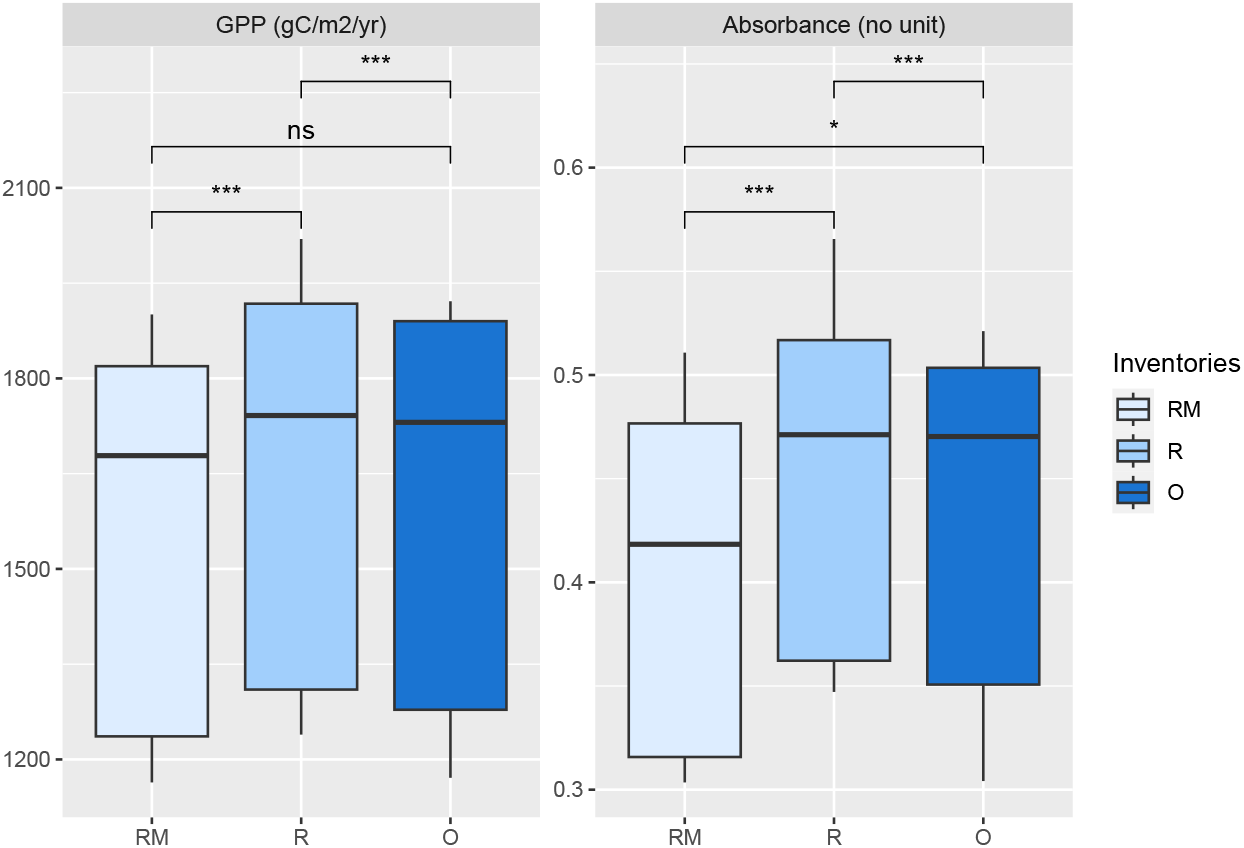
Gross primary production (GPP) and canopy absorbance simulated by PDG-Arena for 13 mixed stands. Three types of inventories were used: regularized monospecific inventories (RM), regularized inventories with species interactions (R) and original inventories (O). Two-sided Wilcoxon signed rank test was used (**: p-value < 0.01, ***: p-value < 0.001).

The structure effect (evaluated by comparing O and R inventories on all 39 stands, not shown here) decreased the GPP by 3.7% (1603 vs 1665 gC/m2/year; p-value < 0.001) and the canopy absorbance by 5.2% (0.428 vs 0.452; p-value < 0.001). Transpiration showed a decrease of 3.2% (226 vs 234 mm; p-value < 0.001) and maximum water shortage a decrease of 1.9% (90.8 vs 92.51 mm; p-value < 0.05).

## 4. Discussion

Given the paucity of forest growth models simulating ecophysiological processes at the individual scale, we developed the individual-based model PDG-Arena from the stand-scale model CASTANEA in order to simulate the carbon, water, and radiation dynamics of mixed forests. PDG-Arena was built with the idea of observing and understanding the properties that emerge in multispecific stands, by integrating tree-level competition and without assuming the presence of positive interactions between heterospecific trees. It uses on the one hand a physiological model parameterized for monospecific stands and on the other hand an individual scale structure that allows trees to interact - the interaction being more or less competitive depending on the functional traits of the individuals and species.

We showed that PDG-Arena was able to reproduce the behavior of CASTANEA when simulating regularized inventories with no species interactions. Thus, the increase in complexity of PDG-Arena, required in order to simulate the functioning and interactions of distinct trees, was not at the cost of decreased performance at stand scale. Even when using original inventories (i.e., integrating the diversity in structure and species), the stand scale results of PDG-Arena were highly correlated to those of CASTANEA. This is explained by the fact that both models are based on LAI, which remains identical for each stand between simulations. Still, PDG-Arena, in comparison to CASTANEA, showed better performance when compared to measurements, in particular on beech (r^2^ +15.4 percentage points) and mixed stands (r^2^ +4.2 percentage points). As shown by the simulations using different types of inventories, the improvement in simulating stand growth is largely explained by the use of original stand structures, letting PDG-Arena simulate the growth of trees of various sizes.

At the individual scale, PDG-Arena explained half of the variability of tree growth, showing that it can capture the competitive status of each tree based on their leaf surface, height and position. However, the mean absolute error was often large and systematic, indicating that the model lacks calibration for each site.

Interestingly, a positive and significant net mixing effect was observed in PDG-Arena simulations on gross primary productivity by comparing simulations with interacting species to equivalent simulations with species in isolation. The simulated overyielding can be attributed to an improvement of canopy absorbance due to species mixing (Figure 4). LAI being equal between each inventory for the same stand, the increased light absorption is hence explained by a greater occupation of the aerial space due to species interactions. This effect, known as canopy packing, has been observed on a variety of mixed forests across Europe (Jucker et al., 2015; Pretzsch, 2019). Canopy packing is commonly decomposed into two mechanism: the phenotypic plasticity of the shape and size of crowns and the vertical stratification (i.e., the occupation by crowns of different vertical strata). Although it is likely to play a role in the functioning of mixed stands (Pretzsch, 2019; Dieler and Pretzsch, 2013), phenotypic plasticity is not yet implemented in PDG-Arena. Thus, our model can only simulate the vertical stratification of crowns, but not their morphological adaptation to their local competitor (see, for example, Jonard et al., 2020 and Morin et al., 2021), potentially leading to an underestimation of overyielding.

The observed overyielding in the French National Forest Inventory for beech-fir mixtures (20%, Toïgo et al., 2015) is greater than the one we simulated. In addition to canopy packing, the real-life overyielding in mixed stands can also be explained by reduced competition for nutrients. Indeed, nutrient content in above-ground biomass and the nitrogen concentration of leaves are likely to be increased by species mixing (Richards et al., 2010). However, competition for nutrients was not integrated in PDG-Arena since its main objective was to build an individual-based model upon the physiological processes that already exist in CASTANEA.

In addition, species mixing increased the yearly water shortage due to increased transpiration (Figure C.8) at equivalent LAI. This confirms the idea that the nature of the diversity-functioning relationship in forests strongly depends on the considered resources (Forrester, 2014). According to our simulations, promoting diverse stands could maximize light interception and growth but would also increase transpiration, which would be detrimental in sites with limited water reserves. In reality, an increase in water use in mixed stands could be counter-balanced by a reduced competition for water between trees of different species (Schume et al., 2004). Although an interspecific differentiation between the water uptake depth has been observed for some species (Schwendenmann et al., 2015), our model cannot simulate this mechanism yet. A comprehensive knowledge of each species water uptake depth is still in construction but could be integrated in process-based models in the near future (Bachofen et al., 2024). Concerning the horizontal distance of tree water uptake, little data exist at the moment. The assumption of a horizontally homogeneous water uptake in our model is justified by the small surface area of the simulated plot.

One limit of this study was the nature of the data used to evaluate the model. Tree growth is an integrative measure that results from carbon, water and light uptake, whereas CASTANEA is calibrated using CO_2_ fluxes (Dufrêne et al., 2005). Moreover, the modeling of carbon allocation, which plays a decisive role in simulating wood growth, is a potential source of error (Davi et al., 2009; Merganičová et al., 2019). Additionally, climate was parameterized at the site scale using an 8 km resolution data set instead of at the stand scale, although climatic variables such as precipitation could vary between stands due to local topography.

## 5. Conclusion

The new individual-based model PDG-Arena we developed is able to simulate the interactions between trees in monospecific and mixed stands and predict their productivity based on an explicit tree inventory. Compared to CASTANEA, PDG-Arena showed improved predictive capability for beech and mixed beech-fir forests. The model can simulate the growth of small-sized stands (less than 1 ha), of regular or irregular structure, and composed of trees of similar or different species (given that the species ecophysiological properties are parametrized in CASTANEA). As PDG-Arena simulates the competition for water and light between trees with no preconceived ideas about the direction of interspecific interaction (from competition to complementarity), it can be used to test specific hypotheses about mixed forests and better understand the diversity-functioning relationship in forests under contrasted scenarios. For example, the model could be used to explore the following open questions, keeping in mind that the answers are largely species-specific and environment-dependent (Ratcliffe et al., 2015; Forrester et al., 2017a): is overyielding more likely to occur in less productive sites (Toïgo et al., 2015)? Can overyielding increase water stress in mixed stands (Forrester et al., 2016)? Are mixed stands more resilient to drought (Grossiord, 2018)? Lastly, being built on the basis of a physio-demo-genetics model, PDG-Arena is suitable to evaluate the evolutionary dynamics of functional traits of a population under various biotic (stand composition, density and structure) and abiotic (soil, climate) constraints, as intraspecific diversity is a major adaptive force in natural tree populations (Lefèvre et al., 2014; Oddou-Muratorio et al., 2020; Fady et al., 2020).

## 6. Declarations

### 6.1. Author contributions

**Camille Rouet**: Conceptualization, Methodology, Software, Visualization, Writing - Original Draft. **Hendrik Davi**: Conceptualization, Supervision. **Arsène Druel**: Methodology, Writing - Review. **Bruno Fady**: Project administration, Supervision, Writing - Review. **Xavier Morin**: Methodology, Data Curation, Supervision, Writing - Review.

### 6.4. Credits

Figures 1 and 2 were designed using images from flaticon.com.

### 6.5. License

For the purpose of Open Access, the authors have applied a CC BY-NC 4.0 public copyright licence to any Author Accepted Manuscript (AAM) version arising from this submission.

## Appendix A. Height-diameter relationship

For each group of trees of the same species and site, a linear model (Equation A.1) was fitted on the logarithms of their measured height (in m) and DBH (in cm) as shown in Figure A.5. The slope and intercept parameter *a* and *b* as well as the coefficients of determination r^2^ are shown in Table A.5 for each group.

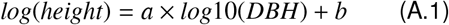

## Appendix B. Supplementary description of PDG-Arena

### Appendix B.1. Computing Leaf Mass per Area

The Leaf Mass per Area (LMA) is a leaf-level trait defined as the mass per unit area of leaves (g/m^2^). LMA varies both in time during leaf growth and in space: leaf mass gain is indeed favored by local irradiance, resulting in an exponentially decreasing distribution of LMA across the canopy from top to bottom. This section describes how the spatial variation of LMA is accounted in PDG-Arena.

In the CASTANEA model, which assumes that the stand is homogeneous and monospecific, the LMA follows a exponentially decreasing function (Davi et al., 2008a):

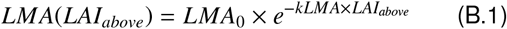

*LAI*_*above*_ is the Leaf Area Index that accounts only for the leaves that are above the considered leaf. *LMA*_0_ and *kLMA* depend on the species and describe the decrease in LMA within the canopy, which is related to the decrease in light intensity within the canopy. Then, the average *LMA* within a layer is obtained by integrating *LMA*(*LAI*_*above*_) within the layer’s vertical boundaries.

In the case of PDG-Arena, the canopy is more structurally complex than in CASTANEA and can include several species. The LMA at a given position of a tree is defined taking all trees into account and using the same formula as in Equation B.1. *LAI*_*above*_ is computed by counting only the leaves of the canopy that are located above the considered leaf. It should be noted that the model is not completely accurate given that the parameter *kLMA* and *LMA*_0_ are those of the species of the considered leaf, although the leaves taken into account in *LAI*_*above*_ potentially come from another species. However, this method does represent the phenomenon of light attenuation which is specific to each individual.

### Appendix B.2. Estimation of the attenuation coefficient with reverse-engineering

In order to know the value of the attenuation coefficients of each species in PDG-Arena, a preliminary simulation is carried out following the CASTANEA model to take advantage of SAIL, its radiation sub-model (Dufrêne et al., 2005). The preliminary simulation is performed for each species on a monospecific and regularized inventory (RM inventory, see section 2.3). We define the attenuation coefficient *k*_1_ at a given time as a function of the incident energy *I*_0_, the energy transmitted by the vegetation *I*_*t*_, and the Leaf Area Index *LAI*, following a Beer-Lambert model:

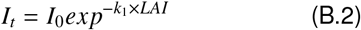

which is equivalent to:

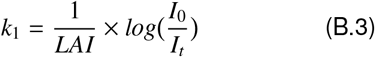

where *I*_*t*_ is defined at any time as the difference between the incident energy and the energy absorbed by the vegetation.

The coefficient of attenuation which is used in SamsaraLight, denoted *k*_2_, is not of the same nature as *k*_1_. Indeed, in Equation B.2, we multiply *k*_1_ by the *LAI* (considering an infinite, horizontally homogeneous, leaf layer) while SamsaraLight multiplies *k*_2_ to the Leaf Area Density *LAD* and the beam path length within a finite, volumetric crown (see Equation 4). Then, to go from one to the other, we must multiply *k*_1_ by *sin*(*β*) (with *β* the angle of height of the sun):

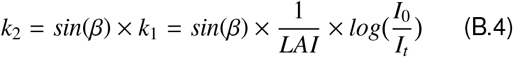

The coefficient *k*_2_ depends on the height of the sun, but also on the frequency domain of the radiation. Indeed, the attenuation coefficient takes into account both the extinction of the rays (defined by the leaf and crown geometry) and the absorption by the leaves which depends on the light frequency. In the following calculations, we distinguish the PAR (photosynthetically active radiation) domain and the NIR (near infrared radiation) domain. It is assumed that these two domains represent the bulk of the incident radiation. To sum up, the attenuation coefficient depends on the species (leaf angle distribution and absorbance rate), the type of radiation (PAR/NIR, direct/diffuse) and the height angle (*β*).

Based on the results of the preliminary CASTANEA simulation, which executes a radiation balance using the SAIL model, we can infer the value of the attenuation coefficients of the plot for direct and diffuse radiations using Equation B.4. In the preliminary simulation, we know for direct rays the value of the height angle *β* at any hour. For diffuse rays, by definition *β* takes every value between 0 and *π/*2 at any hour, so we can’t use the height angle information.

#### Direct Rays

For direct radiation, we estimate an attenuation coefficient for each species by discriminating the PAR and NIR and defining 20 classes of attenuation coefficients corresponding to classes of the height angle *β*, equally distributed between 0 and *π/*2. For each class *i* of *β*, we performed an average on the attenuation coefficients observed during the preliminary simulation for direct radiations:

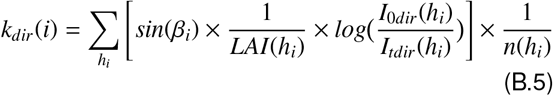

where *k*_*dir*_(*i*) is the mean attenuation coefficient computed from the preliminary simulation results, for direct radiation of the height angle class *i* (which includes *n*(*h*_*i*_) hours). For a given hour of the year *h*_*i*_, *LAI*(*h*_*i*_) is the daily Leaf Area Index of the plot, *I*_0*dir*_(*h*_*i*_), is the incident direct energy and *I*_*tdir*_(*h*_*i*_) is the direct energy transmitted through the canopy.

#### Diffuse Radiation

For diffuse radiation, we discriminate the attenuation coefficient according to the species and radiation domain only. The attenuation coefficient for diffuse light *k*_*di f*_ is assumed to be constant for any sun height angle. To switch from one formulation of the Beer-Lambert law to the other (Equation B.4), a value of *β* is nevertheless needed. Considering that the distribution of diffuse rays along the *β* height angles is uniform, we simplify the equation by using an average computation. Then, we use 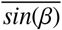, the average of *sin*(*β*) for *β* going from 0 to *π/*2 (which is about 0.637). For a species and a radiative domain, we compute an average on every day of year of the observed attenuation coefficient during the preliminary simulation:

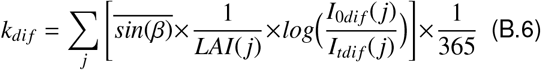

with, for day *j, LAI*(*j*) the Leaf Area Index, *I*_0*di f*_ (*j*) the incident diffuse energy and *I*_*tdi f*_ (*j*) the diffuse energy transmitted through canopy.

### Appendix B.3. Distribution of radiations into canopy layers and into sun and shade leaves

In CASTANEA, the energy absorbed by the canopy is distributed into five layers of leaves, which are themselves divided into leaves in direct light (called sun leaves) and leaves in the shade. We present here how PDG-Arena operates the distribution of the absorbed energy by individual crowns.

#### Proportion of sun leaves of a tree

The proportion of sun leaves of a crown, i.e., of its leaves subjected to direct radiation, is given by a formula borrowed from the HETEROFOR model (Jonard et al., 2020). Two factors define the shading received by the leaves of a tree: on the one hand, the external shading provided by the competing trees, giving the proportion of sun leaves *pS un*_*ext*_; on the other hand, the internal shading provided by the own leaves of a tree, giving the proportion of sun leaves *pS un*_*int*_.

The shading provided by the competitors is given by the ratio of the direct incident energy above the tree *I*_*d*0_(*aboveTree*) to the potential direct incident energy *I*_*d*0_(*potential*), which is computed by SamsaraLight by ignoring neighbors trees:

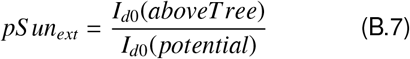

The second quotient to be evaluated is the proportion of the leaves of the tree shaded by its own leaves. The shading by the leaves of the tree itself follows the same relationship as the direct radiation within the tree, that is to say a Beer-Lambert law:

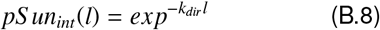

where *pS un*_*int*_(*l*) is the proportion of sun leaves remaining after the radiation passes through the crown, with *l* the cumulative LAI encountered by the passing beam and *k*_*dir*_ the tree extinction coefficient for direct PAR. The proportion of sun leaves at the crown entrance is supposed to be 1, ignoring leaves shaded by neighboring trees.

We can compute *LAI*_*sun*−*int*_, the amount of leaves that are not shaded by leaves of the same tree. To do this, we need to integrate *pS un*_*int*_(*l*) for *l* ranging from 0 to *LAI*, the Leaf Area Index of the tree:

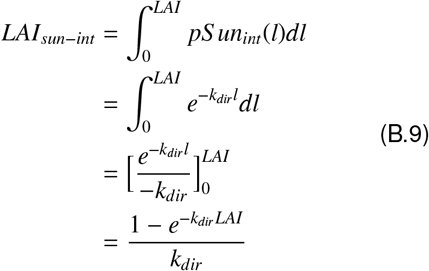

Thus, *pS un*_*int*_ = *LAI*_*sun*−*int*_*/LAI* represents the proportion of leaf remaining in the light when shaded by the tree’s own leaves.

Finally, the proportion of sun leaves of a tree is *pS un*_*tree*_ = *pS un*_*ext*_ × *pS un*_*int*_.

#### Distribution of radiations by layer

If SamsaraLight allows us to know the amount of energy absorbed per tree according to each domain (PAR/NIR) and type of energy (direct/diffused), noted *E*_*tree*_, it does not allow us to distribute this amount between layers, differentiating leaves with high interception and leaves with low interception. To do so, we firstly divide the leaf surface of a tree into *n* equal-sized layers, and we assume that the radiative characteristics are homogeneous within a layer. We define a distribution function *f*_*i*_, that determines *E*_*i*_, the amount of energy that is absorbed by layer *i*:

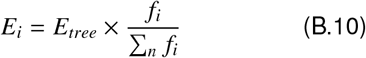

We assume that the distribution *f*_*i*_ is affected by the light interception from leaf surface that is located above the layer (whether it belongs to other trees or to the same tree). Then, we define a simple stand-scale model that describes the level of energy transmitted through the stand using the Beer-Lambert law. At any level of height located under a quantity of leaves *LAI*_*above*_, the proportion of light transmitted through these leaves is:

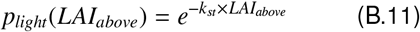

with *k*_*st*_ the stand level attenuation coefficient.

*LAI*_*above*_ is calculated by counting the amount of leaves above the leaf layer under consideration, knowing the position and shape of each individual. A homogeneous distribution of leaf density within each individual crown is assumed. We do not consider the plot slope in this calculation, i.e., only tree height defines whether the leaves of one tree are higher than those of another.

Finally, to calculate *f*_*i*_, the fraction of energy absorbed by any layer *i* of a crown, we compute the average value of *p*_*light*_ inside the layer by integrating it within its boundaries *LAI*_*above*_(*i* − 1) and *LAI*_*above*_(*i*):

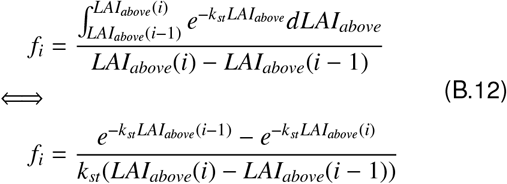

The proportion *f*_*i*_ is computed for each type of radiation (direct/diffuse and PAR/NIR).

### Appendix B.4. Reduction of absorbed radiations in SamsaraLight

In SamsaraLight standard mode, the foliage is assumed to be at its maximum during the whole process. Thus, the energy absorbed by the trees when their leaf area is in reality lower must be revised downwards, especially for deciduous trees, which lose all their leaves in autumn. For each individual, a ratio depending on its LAI is computed each day to represent the evolution of its absorption level from 0 to 1. The level of absorption is supposed to follow the dynamic of the Beer-Lambert law:

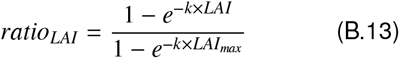

For each radiation domain, *k* is the attenuation coefficient of a tree and *ratio*_*LAI*_ is applied to its absorbed energy to take off the surplus. Nevertheless, the removed energy must be redistributed, because if it had not been intercepted, this energy would have been distributed among the other absorbing elements (crowns or soil cells). At this point, it is no longer possible to know to which element the energy should be distributed. Then, the extracted energy is redistributed to all absorbing elements, proportionally to their level of absorbed energy (after reduction according to LAI), which represents their relative interception capacity.

## Appendix C. Supplementary results

Figure C.6 shows the simulated versus measured wood volume increment per stand for the 39 stands (including the outliers) using the CASTANEA model (with RM inventories) and the PDG-Arena model (with O inventories). Table C.6 shows the performance of the models at stand scale based on the r^2^ and MAPE coefficients, computed without discarding the two silver fir outlier stands.

Figure C.7 shows the simulated versus measured wood volume increment per tree for the 37 stands using the PDG-Arena model with O inventories. Table C.7 shows the individual-scale performances in terms of r^2^ and MAPE.

Figure C.8 shows the maximum water shortage during an average year (i.e., the maximum difference reached during a year between the current and full useful reserve, in mm) and yearly transpiration simulated by PDG-Arena for 13 mixed stands using RM, R and O inventories.

**Figure A.5:**
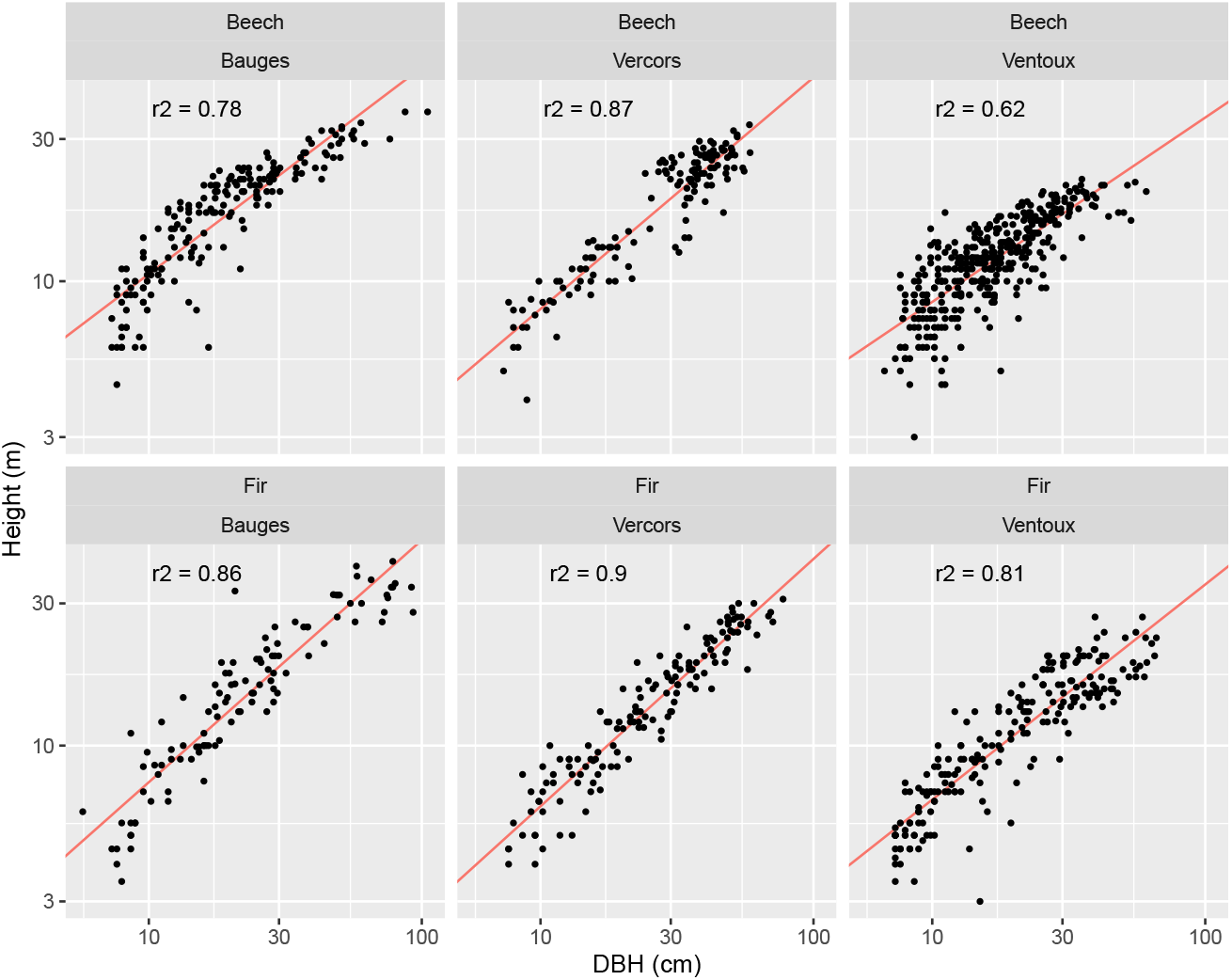
Relationship between measured height and DBH. The red line indicates the model fitted on logarithmic values.

**Table A.5:** Parameters of the height-DBH model described in Equation A.1.

| Site | Species | $a$ | $b$ | $r^2$ |
| --- | --- | --- | --- | --- |
| Bauges | Beech | 0.69 | 0.33 | 0.78 |
| Bauges | Fir | 0.81 | 0.065 | 0.86 |
| Ventoux | Beech | 0.62 | 0.31 | 0.62 |
| Ventoux | Fir | 0.72 | 0.097 | 0.81 |
| Vercors | Beech | 0.78 | 0.13 | 0.87 |
| Vercors | Fir | 0.83 | 0.033 | 0.90 |

**Figure C.6:**
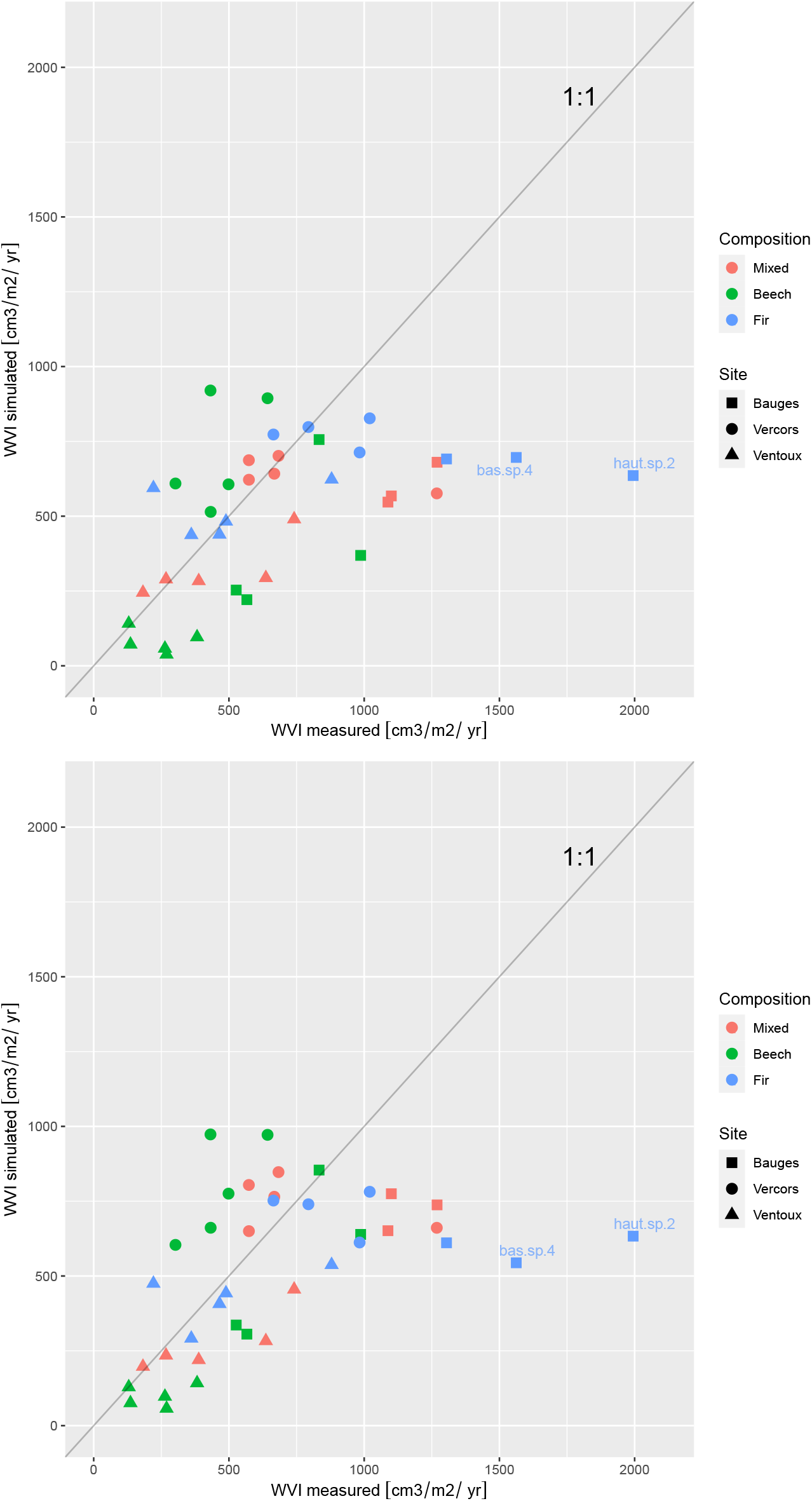
Simulated versus measured wood volume increment for the 39 stands using the CASTANEA model and RM inventories (top) and using the PDG-Arena model and original inventories (O) (bottom). Labelled points are the outlier stands.

**Table C.6:** Evaluation of the performances of PDG-Arena and CASTANEA without discarding outliers. Coefficient of determination (r^2^, in %) and Mean Absolute Percentage Error (MAPE, in %) were computed for the simulated versus measured yearly wood volume increment per stand over the period 1996-2013. Inventories are characterized as: ‘RM’ (regularized and monospecific, i.e., without species interactions), ’R’ (regularized, but with species interactions) and ’O’ (original inventories).

| Set | Model | Inventories | $r^2$ | MAPE |
| --- | --- | --- | --- | --- |
| All stands | CASTANEA | RM | 25.7 | 41.7 |
|  | PDG-Arena | RM | 26.5 | 41.6 |
|  | PDG-Arena | R | 26.4 | 42.8 |
|  | PDG-Arena | O | 24.0 | 41.7 |
| Mixed | CASTANEA | RM | 36.3 | 30.1 |
|  | PDG-Arena | RM | 37.6 | 30.7 |
|  | PDG-Arena | R | 36.3 | 33.1 |
|  | PDG-Arena | O | 40.5 | 31.5 |
| Beech pure | CASTANEA | RM | 22.9 | 55.3 |
|  | PDG-Arena | RM | 25.0 | 57.4 |
|  | PDG-Arena | R | 24.7 | 57.9 |
|  | PDG-Arena | O | 38.3 | 53.9 |
| Fir pure | CASTANEA | RM | 18.0 | 38.4 |
|  | PDG-Arena | RM | 24.8 | 34.9 |
|  | PDG-Arena | R | 23.7 | 35.6 |
|  | PDG-Arena | O | 19.1 | 38.6 |

**Table C.7:** Performance at the individual scale of the PDG-Arena model using original inventories (O). r^2^ and MAPE, expressed in %, were computed on set of trees of the same site, type of stand and species.

| Site | Stand type | Species | $r^2$ | MAPE |
| --- | --- | --- | --- | --- |
| Bauges | Mixed | Beech | 36 | 70 |
| Bauges | Mixed | Fir | 62 | 68 |
| Bauges | Pure beech | Beech | 64 | 63 |
| Bauges | Pure fir | Fir | 20 | 73 |
| Ventoux | Mixed | Beech | 40 | 95 |
| Ventoux | Mixed | Fir | 59 | 50 |
| Ventoux | Pure beech | Beech | 40 | 69 |
| Ventoux | Pure fir | Fir | 43 | 95 |
| Vercors | Mixed | Beech | 51 | 146 |
| Vercors | Mixed | Fir | 49 | 68 |
| Vercors | Pure beech | Beech | 51 | 115 |
| Vercors | Pure fir | Fir | 48 | 67 |

**Figure C.7:**
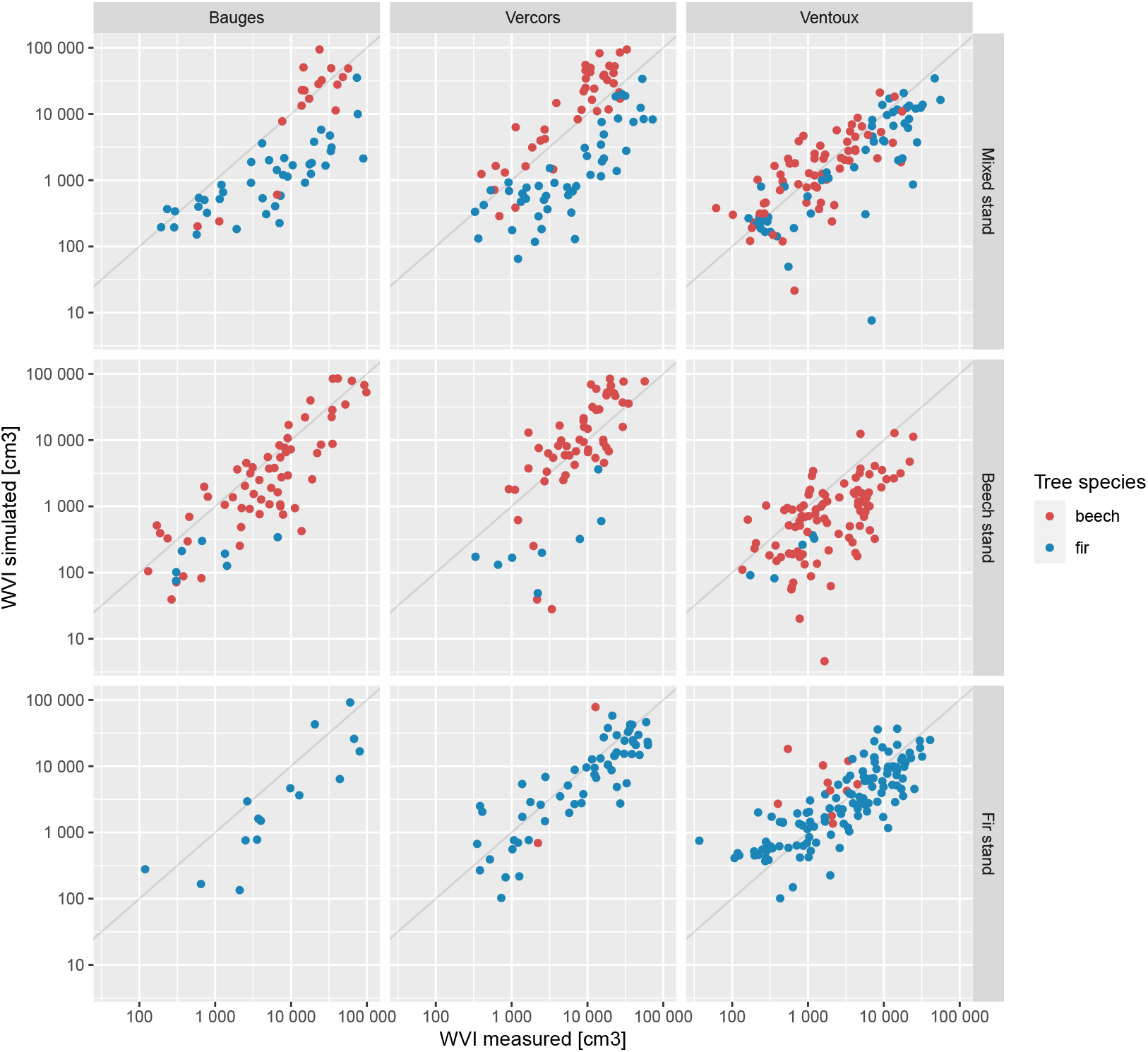
Simulated versus measured wood volume increment for every cored trees using the PDG-Arena model and original inventories (log scale).

**Figure C.8:**
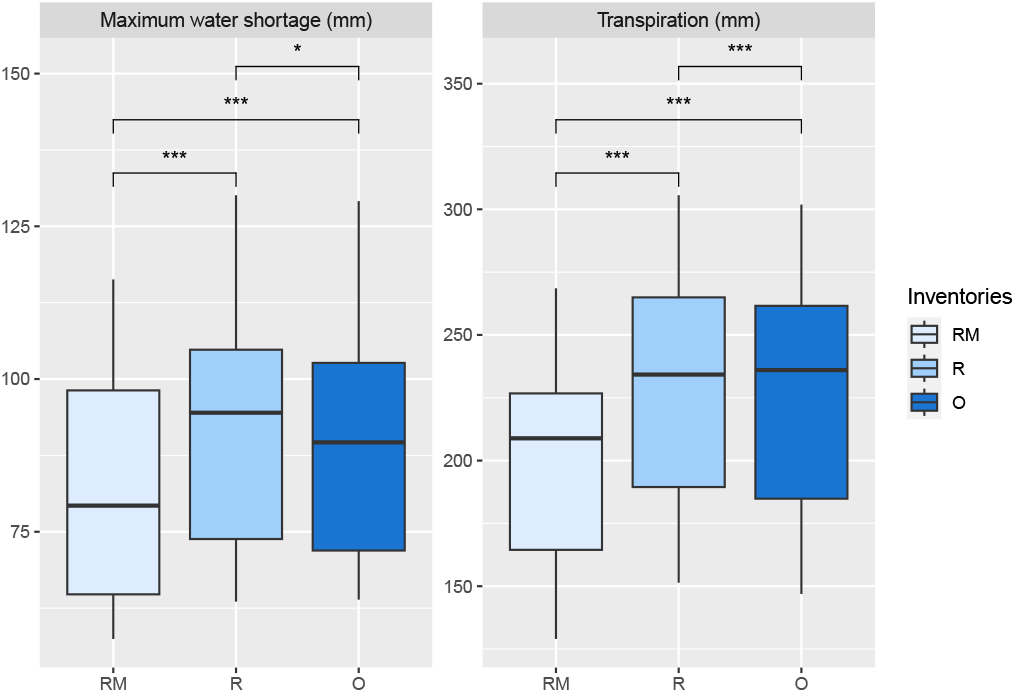
Maximum water shortage (defined as the yearly maximum difference reached between the current and full useful reserve) and yearly transpiration simulated by PDG-Arena for 13 mixed stands.Three types of inventories were used: regularized monospecific inventories (RM), regularized inventories with species interactions (R) and original inventories (O). Two-sided Wilcoxon signed rank test was used (*: p-value < 0.05, ***: p-value < 0.001).

## References

Ammer, C., 2019. Diversity and forest productivity in a changing climate. New Phytologist 221, 50–66. doi:10.1111/nph.15263.

González de Andrés, E., 2019. Interactions between Climate and Nutrient Cycles on Forest Response to Global Change: The Role of Mixed Forests. Forests 10, 609. doi:10.3390/f10080609.

Bachofen, C., Tumber-Dávila, S.J., Mackay, D.S., McDowell, N.G., Carminati, A., Klein, T., Stocker, B.D., Mencuccini, M., Grossiord, C., 2024. Tree water uptake patterns across the globe. New Phytologist 242, 1891–1910. doi:10.1111/nph.19762.

Barbosa, L.O., dos Santos, J.A., Gonçalves, A.F.A., Campoe, O.C., Scolforo, J.R.S., Scolforo, H.F., 2023. Competition in forest plantations: Empirical and process-based modelling in pine and eucalypt plantations. Ecological Modelling 483, 110410. doi:10.1016/j.ecolmodel.2023.110410.

Baret, F., Weiss, M., Lacaze, R., Camacho, F., Makhmara, H., Pacholcyzk, P., Smets, B., 2013. GEOV1: LAI and FAPAR essential climate variables and FCOVER global time series capitalizing over existing products. Part1: Principles of development and production. Remote Sensing of Environment 137, 299–309. doi:10.1016/j.rse.2012.12.027.

Bauhus, J., Forrester, D.I., Pretzsch, H., 2017. From Observations to Evidence About Effects of Mixed-Species Stands, in: Pretzsch, H., Forrester, D.I., Bauhus, J. (Eds.), Mixed-Species Forests: Ecology and Management. Springer, Berlin, Heidelberg, pp. 27–71. doi:10.1007/978-3-662-54553-9_2.

Bonan, G.B., 2008. Forests and Climate Change: Forcings, Feedbacks, and the Climate Benefits of Forests. Science 320, 1444–1449. doi:10.1126/science.1155121.

Cordonnier, T., Smadi, C., Kunstler, G., Courbaud, B., 2019. Asymmetric competition, ontogenetic growth and size inequality drive the difference in productivity between two-strata and one-stratum forest stands. Theoretical Population Biology 130, 83–93. doi:10.1016/j.tpb.2019.07.001.

Courbaud, B., de Coligny, F., Cordonnier, T., 2003. Simulating radiation distribution in a heterogeneous Norway spruce forest on a slope. Agricultural and Forest Meteorology 116, 1–18. doi:10.1016/S0168-1923(02)00254-X.

Cuddington, K., Fortin, M.J., Gerber, L.R., Hastings, A., Liebhold, A., O’Connor, M., Ray, C., 2013. Process-based models are required to manage ecological systems in a changing world. Ecosphere 4, art20. doi:10.1890/ES12-00178.1.

Davi, H., Barbaroux, C., Dufrêne, E., François, C., Montpied, P., Bréda, N., Badeck, F., 2008a. Modelling leaf mass per area in forest canopy as affected by prevailing radiation conditions. Ecological Modelling 211, 339–349. doi:10.1016/j.ecolmodel.2007.09.012.

Davi, H., Barbaroux, C., Francois, C., Dufrêne, E., 2009. The fundamental role of reserves and hydraulic constraints in predicting LAI and carbon allocation in forests. Agricultural and Forest Meteorology 149, 349–361. doi:10.1016/j.agrformet.2008.08.014.

Davi, H., Baret, F., Huc, R., Dufrêne, E., 2008b. Effect of thinning on LAI variance in heterogeneous forests. Forest Ecology and Management 256, 890–899. doi:10.1016/j.foreco.2008.05.047.

Davi, H., Cailleret, M., 2017. Assessing drought-driven mortality trees with physiological process-based models. Agricultural and Forest Meteorology 232, 279–290. doi:10.1016/j.agrformet.2016.08.019.

Deleuze, C., Morneau, F., Renaud, J., Vivien, Y., Rivoire, M., Santenoise, P., Longuetaud, F., Mothe, F., Hervé, J., Vallet, P., 2014. Estimer le volume total d’un arbre, quelles que soient l’essence, la taille, la sylviculture, la station. Rendez-vous Techniques de l’ONF, 22–32.

Dieler, J., Pretzsch, H., 2013. Morphological plasticity of European beech (Fagus sylvatica L.) in pure and mixed-species stands. Forest Ecology and Management 295, 97–108. doi:10.1016/j.foreco.2012.12.049.

Dufour-Kowalski, S., Courbaud, B., Dreyfus, P., Meredieu, C., de Coligny, F., 2012. Capsis: an open software framework and community for forest growth modelling. Annals of Forest Science 69, 221–233. doi:10.1007/s13595-011-0140-9.

Dufrêne, E., Davi, H., François, C., Maire, G.l., Dantec, V.L., Granier, A., 2005. Modelling carbon and water cycles in a beech forest. Part I: Model description and uncertainty analysis on modelled NEE. Ecological Modelling 185, 407–436. doi:10/fjnfgr.

Dă nescu, A., Albrecht, A.T., Bauhus, J., 2016. Structural diversity promotes productivity of mixed, uneven-aged forests in southwestern Germany. Oecologia 182, 319–333. doi:10.1007/s00442-016-3623-4.

Fady, B., Aravanopoulos, F., Benavides, R., González-Martínez, S., Grivet, D., Lascoux, M., Lindner, M., Rellstab, C., Valladares, F., Vinceti, B., 2020. Genetics to the rescue: managing forests sustainably in a changing world. Tree Genetics & Genomes 16, 80. doi:10.1007/s11295-020-01474-8.

Fatichi, S., Leuzinger, S., Körner, C., 2014. Moving beyond photosynthesis: from carbon source to sink-driven vegetation modeling. New Phytologist 201, 1086–1095. doi:10.1111/nph.12614.

Fontes, L., Bontemps, J.D., Bugmann, H., Oijen, M.v., Gracia, C., Kramer, K., Lindner, M., Rötzer, T., Skovsgaard, J.P., 2010. Models for supporting forest management in a changing environment. Forest Systems 19, 8–29.

Forrester, D.I., 2014. The spatial and temporal dynamics of species interactions in mixed-species forests: From pattern to process. Forest Ecology and Management 312, 282–292. doi:10.1016/j.foreco.2013.10.003.

Forrester, D.I., Ammer, C., Annighöfer, P.J., Avdagic, A., Barbeito, I., Bielak, K., Brazaitis, G., Coll, L., del Río, M., Drössler, L., Heym, M., Hurt, V., Löf, M., Matović, B., Meloni, F., den Ouden, J., Pach, M., Pereira, M.G., Ponette, Q., Pretzsch, H., Skrzyszewski, J., Stojanović, D., Svoboda, M., Ruiz-Peinado, R., Vacchiano, G., Verheyen, K., Zlatanov, T., Bravo-Oviedo, A., 2017a. Predicting the spatial and temporal dynamics of species interactions in Fagus sylvatica and Pinus sylvestris forests across Europe. Forest Ecology and Management 405, 112–133. doi:10.1016/j.foreco.2017.09.029.

Forrester, D.I., Bauhus, J., 2016. A Review of Processes Behind Diversity—Productivity Relationships in Forests. Current Forestry Reports 2, 45–61. doi:10.1007/s40725-016-0031-2.

Forrester, D.I., Bonal, D., Dawud, S., Gessler, A., Granier, A., Pollastrini, M., Grossiord, C., 2016. Drought responses by individual tree species are not often correlated with tree species diversity in European forests. Journal of Applied Ecology 53, 1725–1734. doi:10.1111/1365-2664.12745.

Forrester, D.I., Tachauer, I.H.H., Annighoefer, P., Barbeito, I., Pretzsch, H., Ruiz-Peinado, R., Stark, H., Vacchiano, G., Zlatanov, T., Chakraborty, T., Saha, S., Sileshi, G.W., 2017b. Generalized biomass and leaf area allometric equations for European tree species incorporating stand structure, tree age and climate. Forest Ecology and Management 396, 160–175. doi:10.1016/j.foreco.2017.04.011.

Gonçalves, A.F.A., Santos, J.A.d., França, L.C.d.J., Campoe, O.C., Altoé, T.F., Scolforo, J.R.S., 2021. Use of the process-based models in forest research: a bibliometric review. CERNE 27, e. doi:10.1590/01047760202127012769.

Granier, A., Bréda, N., Biron, P., Villette, S., 1999. A lumped water balance model to evaluate duration and intensity of drought constraints in forest stands. Ecological Modelling 116, 269–283. doi:10.1016/S0304-3800(98)00205-1.

Grossiord, C., 2018. Having the right neighbors: how tree species diversity modulates drought impacts on forests. New Phytologist 228, 42–49. doi:10.1111/nph.15667.

Grossiord, C., Granier, A., Ratcliffe, S., Bouriaud, O., Bruelheide, H., Chećko, E., Forrester, D.I., Dawud, S.M., Finér, L., Pollastrini, M., Scherer-Lorenzen, M., Valladares, F., Bonal, D., Gessler, A., 2014. Tree diversity does not always improve resistance of forest ecosystems to drought. Proceedings of the National Academy of Sciences 111, 14812–14815. doi:10.1073/pnas.1411970111.

Guillemot, J., Kunz, M., Schnabel, F., Fichtner, A., Madsen, C.P., Gebauer, T., Härdtle, W., von Oheimb, G., Potvin, C., 2020. Neighbourhood-mediated shifts in tree biomass allocation drive overyielding in tropical species mixtures. New Phytologist 228, 1256–1268. doi:10.1111/nph.16722.

Jonard, M., André, F., de Coligny, F., de Wergifosse, L., Beudez, N., Davi, H., Ligot, G., Ponette, Q., Vincke, C., 2020. HETEROFOR 1.0: a spatially explicit model for exploring the response of structurally complex forests to uncertain future conditions – Part 1: Carbon fluxes and tree dimensional growth. Geoscientific Model Development 13, 905–935. doi:10.5194/gmd-13-905-2020.

Jourdan, M., Cordonnier, T., Dreyfus, P., Riond, C., de Coligny, F., Morin, X., 2021. Managing mixed stands can mitigate severe climate change impacts on French alpine forests. Regional Environmental Change 21, 78. doi:10.1007/s10113-021-01805-y.

Jourdan, M., Kunstler, G., Morin, X., 2020. How neighbourhood interactions control the temporal stability and resilience to drought of trees in mountain forests. Journal of Ecology 108, 666–677. doi:10.1111/1365-2745.13294.

Jourdan, M., Lebourgeois, F., Morin, X., 2019. The effect of tree diversity on the resistance and recovery of forest stands in the French Alps may depend on species differences in hydraulic features. Forest Ecology and Management 450, 117486. doi:10.1016/j.foreco.2019.117486.

Jucker, T., Bouriaud, O., Avacaritei, D., Dănilă, I., Duduman, G., Valladares, F., Coomes, D.A., 2014. Competition for light and water play contrasting roles in driving diversity-productivity relationships in Iberian forests. Journal of Ecology 102, 1202–1213. doi:10.1111/1365-2745.12276.

Jucker, T., Bouriaud, O., Coomes, D.A., 2015. Crown plasticity enables trees to optimize canopy packing in mixed-species forests. Functional Ecology 29, 1078–1086. doi:10.1111/1365-2435.12428.

Korzukhin, M.D., Ter-Mikaelian, M.T., Wagner, R.G., 1996. Process versus empirical models: which approach for forest ecosystem management? Canadian Journal of Forest Research 26, 879–887. doi:10.1139/x26-096.

Lefèvre, F., Boivin, T., Bontemps, A., Courbet, F., Davi, H., DurandGillmann, M., Fady, B., Gauzere, J., Gidoin, C., Karam, M.J., Lalagüe, H., Oddou-Muratorio, S., Pichot, C., 2014. Considering evolutionary processes in adaptive forestry. Annals of Forest Science 71, 723–739. doi:10.1007/s13595-013-0272-1.

Leuning, R., Kelliher, F.M., Pury, D.G.G.D., Schulze, E.D., 1995. Leaf nitrogen, photosynthesis, conductance and transpiration: scaling from leaves to canopies. Plant, Cell & Environment 18, 1183–1200. doi:10.1111/j.1365-3040.1995.tb00628.x.

Liang, J., Crowther, T.W., Picard, N., Wiser, S., Zhou, M., Alberti, G., Schulze, E.D., McGuire, A.D., Bozzato, F., Pretzsch, H., de Miguel, S., Paquette, A., Hérault, B., Scherer-Lorenzen, M., Barrett, C.B., Glick, H.B., Hengeveld, G.M., Nabuurs, G.J., Pfautsch, S., Viana, H., Vibrans, A.C., Ammer, C., Schall, P., Verbyla, D., Tchebakova, N., Fischer, M., Watson, J.V., Chen, H.Y.H., Lei, X., Schelhaas, M.J., Lu, H., Gianelle, D., Parfenova, E.I., Salas, C., Lee, E., Lee, B., Kim, H.S., Bruelheide, H., Coomes, D.A., Piotto, D., Sunderland, T., Schmid, B., Gourlet-Fleury, S., Sonké, B., Tavani, R., Zhu, J., Brandl, S., Vayreda, J., Kitahara, F., Searle, E.B., Neldner, V.J., Ngugi, M.R., Baraloto, C., Frizzera, L., Bałazy, R., Oleksyn, J., Zawiła-Niedźwiecki, T., Bouriaud, O., Bussotti, F., Finér, L., Jaroszewicz, B., Jucker, T., Valladares, F., Jagodzinski, A.M., Peri, P.L., Gonmadje, C., Marthy, W., O’Brien, T., Martin, E.H., Marshall, A.R., Rovero, F., Bitariho, R., Niklaus, P.A., Alvarez-Loayza, P., Chamuya, N., Valencia, R., Mortier, F., Wortel, V., Engone-Obiang, N.L., Ferreira, L.V., Odeke, D.E., Vasquez, R.M., Lewis, S.L., Reich, P.B., 2016. Positive biodiversity-productivity relationship predominant in global forests. Science 354. doi:10.1126/science.aaf8957.

Lindner, M., Maroschek, M., Netherer, S., Kremer, A., Barbati, A., Garcia-Gonzalo, J., Seidl, R., Delzon, S., Corona, P., Kolström, M., Lexer, M.J., Marchetti, M., 2010. Climate change impacts, adaptive capacity, and vulnerability of European forest ecosystems. Forest Ecology and Management 259, 698–709. doi:10.1016/j.foreco.2009.09.023.

Loreau, M., 2010. CHAPTER 3. Biodiversity and Ecosystem Functioning, in: From Populations to Ecosystems. Princeton University Press, pp. 56–78. doi:10.1515/9781400834167.56.

Mas, E., Cochard, H., Deluigi, J., Didion-Gency, M., Martin-StPaul, N., Morcillo, L., Valladares, F., Vilagrosa, A., Grossiord, C., 2024. Interactions between beech and oak seedlings can modify the effects of hotter droughts and the onset of hydraulic failure. New Phytologist 241, 1021–1034. doi:10.1111/nph.19358.

Merganičová, K., Merganič, J., Lehtonen, A., Vacchiano, G., Sever, M.Z.O., Augustynczik, A.L.D., Grote, R., Kyselová, I., Mäkelä, A., Yousefpour, R., Krejza, J., Collalti, A., Reyer, C.P.O., 2019. Forest carbon allocation modelling under climate change. Tree Physiology 39, 1937–1960. doi:10/ghkr6m.

Messier, C., Bauhus, J., Sousa-Silva, R., Auge, H., Baeten, L., Barsoum, N., Bruelheide, H., Caldwell, B., Cavender-Bares, J., Dhiedt, E., Eisenhauer, N., Ganade, G., Gravel, D., Guillemot, J., Hall, J.S., Hector, A., Hérault, B., Jactel, H., Koricheva, J., Kreft, H., Mereu, S., Muys, B., Nock, C.A., Paquette, A., Parker, J.D., Perring, M.P., Ponette, Q., Potvin, C., Reich, P.B., Scherer-Lorenzen, M., Schnabel, F., Verheyen, K., Weih, M., Wollni, M., Zemp, D.C., 2022. For the sake of resilience and multifunctionality, let’s diversify planted forests! Conservation Letters 15, e12829. doi:10.1111/conl.12829.

Metz, J., Annighöfer, P., Schall, P., Zimmermann, J., Kahl, T., Schulze, E.D., Ammer, C., 2016. Site-adapted admixed tree species reduce drought susceptibility of mature European beech. Global Change Biology 22, 903–920. doi:10.1111/gcb.13113.

Monteith, J., 1965. Evaporation and environment. Symposia of the Society for Experimental Biology.

Morin, X., Bugmann, H., Coligny, F.d., Martin-StPaul, N., Cailleret, M., Limousin, J.M., Ourcival, J.M., Prevosto, B., Simioni, G., Toigo, M., Vennetier, M., Catteau, E., Guillemot, J., 2021. Beyond forest succession: A gap model to study ecosystem functioning and tree community composition under climate change. Functional Ecology n/a. doi:10.1111/1365-2435.13760.

Morin, X., Fahse, L., de Mazancourt, C., Scherer-Lorenzen, M., Bugmann, H., 2014. Temporal stability in forest productivity increases with tree diversity due to asynchrony in species dynamics. Ecology Letters 17, 1526–1535. doi:10.1111/ele.12357.

Morin, X., Fahse, L., Scherer-Lorenzen, M., Bugmann, H., 2011. Tree species richness promotes productivity in temperate forests through strong complementarity between species: Species richness promotes forest productivity. Ecology Letters 14, 1211–1219. doi:10.1111/j.1461-0248.2011.01691.x.

Oddou-Muratorio, S., Davi, H., Lefèvre, F., 2020. Integrating evolutionary, demographic and ecophysiological processes to predict the adaptive dynamics of forest tree populations under global change. Tree Genetics & Genomes 16, 67. doi:10.1007/s11295-020-01451-1.

Oddou-Muratorio, S., Davi, H., 2014. Simulating local adaptation to climate of forest trees with a Physio-Demo-Genetics model. Evolutionary Applications 7, 453–467. doi:10.1111/eva.12143.

Pardos, M., del Río, M., Pretzsch, H., Jactel, H., Bielak, K., Bravo, F., Brazaitis, G., Defossez, E., Engel, M., Godvod, K., Jacobs, K., Jansone, L., Jansons, A., Morin, X., Nothdurft, A., Oreti, L., Ponette, Q., Pach, M., Riofrío, J., Ruíz-Peinado, R., Tomao, A., Uhl, E., Calama, R., 2021. The greater resilience of mixed forests to drought mainly depends on their composition: Analysis along a climate gradient across Europe. Forest Ecology and Management 481, 118687. doi:10.1016/j.foreco.2020.118687.

Piotto, D., 2008. A meta-analysis comparing tree growth in monocultures and mixed plantations. Forest Ecology and Management 255, 781–786. doi:10.1016/j.foreco.2007.09.065.

van der Plas, F., Manning, P., Allan, E., Scherer-Lorenzen, M., Verheyen, K., Wirth, C., Zavala, M.A., Hector, A., Ampoorter, E., Baeten, L., Barbaro, L., Bauhus, J., Benavides, R., Benneter, A., Berthold, F., Bonal, D., Bouriaud, O., Bruelheide, H., Bussotti, F., Carnol, M., Castagneyrol, B., Charbonnier, Y., Coomes, D., Coppi, A., Bastias, C.C., Muhie Dawud, S., De Wandeler, H., Domisch, T., Finér, L., Gessler, A., Granier, A., Grossiord, C., Guyot, V., Hättenschwiler, S., Jactel, H., Jaroszewicz, B., Joly, F.X., Jucker, T., Koricheva, J., Milligan, H., Müller, S., Muys, B., Nguyen, D., Pollastrini, M., Raulund-Rasmussen, K., Selvi, F., Stenlid, J., Valladares, F., Vesterdal, L., Zielínski, D., Fischer, M., 2016. Jack-of-all-trades effects drive biodiversity–ecosystem multifunctionality relationships in European forests. Nature Communications 7, 11109. doi:10.1038/ncomms11109.

Porté, A., Bartelink, H.H., 2002. Modelling mixed forest growth: a review of models for forest management. Ecological Modelling 150, 141–188. doi:10.1016/S0304-3800(01)00476-8.

Pretzsch, H., 2019. The Effect of Tree Crown Allometry on Community Dynamics in Mixed-Species Stands versus Monocultures. A Review and Perspectives for Modeling and Silvicultural Regulation. Forests 10, 810. doi:10.3390/f10090810.

Pretzsch, H., Forrester, D.I., Bauhus, J. (Eds.), 2017. Mixed-species forests : ecology and management. Springer, Berlin.

Pretzsch, H., Forrester, D.I., Rötzer, T., 2015. Representation of species mixing in forest growth models. A review and perspective. Ecological Modelling 313, 276–292. doi:10.1016/j.ecolmodel.2015.06.044.

Pretzsch, H., Schütze, G., Uhl, E., 2013. Resistance of European tree species to drought stress in mixed versus pure forests: evidence of stress release by inter-specific facilitation. Plant Biology 15, 483–495. doi:10.1111/j.1438-8677.2012.00670.x.

Ratcliffe, S., Holzwarth, F., Nadrowski, K., Levick, S., Wirth, C., 2015. Tree neighbourhood matters – Tree species composition drives diversity–productivity patterns in a near-natural beech forest. Forest Ecology and Management 335, 225–234. doi:10.1016/j.foreco.2014.09.032.

Ratcliffe, S., Liebergesell, M., Ruiz-Benito, P., González, J.M., Castañeda, J.M.M., Kändler, G., Lehtonen, A., Dahlgren, J., Kattge, J., Peñuelas, J., Zavala, M.A., Wirth, C., 2016. Modes of functional biodiversity control on tree productivity across the European continent. Global Ecology and Biogeography 25, 251–262. doi:10.1111/geb.12406.

Reyer, C., 2015. Forest Productivity Under Environmental Change—a Review of Stand-Scale Modeling Studies. Current Forestry Reports 1, 53–68. doi:10.1007/s40725-015-0009-5.

Richards, A.E., Forrester, D.I., Bauhus, J., Scherer-Lorenzen, M., 2010. The influence of mixed tree plantations on the nutrition of individual species: a review. Tree Physiology 30, 1192–1208. doi:10.1093/treephys/tpq035.

Rolland, C., 2003. Spatial and Seasonal Variations of Air Temperature Lapse Rates in Alpine Regions. Journal of Climate 16, 1032–1046. doi:10.1175/1520-0442(2003)016<1032:SASVOA>2.0.CO;2.

Rouet, C., Morin, X., Druel, A., 2024. Inputs, results data and analysis script for the evaluation of the PDG-Arena forest growth model on beech-fir stands. doi:10.5281/zenodo.12191049.

del Río, M., Pretzsch, H., Ruiz-Peinado, R., Jactel, H., Coll, L., Löf, M., Aldea, J., Ammer, C., Avdagić, A., Barbeito, I., Bielak, K., Bravo, F., Brazaitis, G., Cerný, J., Collet, C., Condés, S., Drössler, L., Fabrika, M., Heym, M., Holm, S.O., Hylen, G., Jansons, A., Kurylyak, V., Lombardi, F., Matović, B., Metslaid, M., Motta, R., Nord-Larsen, T., Nothdurft, A., den Ouden, J., Pach, M., Pardos, M., Poeydebat, C., Ponette, Q., Pérot, T., Reventlow, D.O.J., Sitko, R., Sramek, V., Steckel, M., Svoboda, M., Verheyen, K., Vospernik, S., Wolff, B., Zlatanov, T., Bravo-Oviedo, A., 2022. Emerging stability of forest productivity by mixing two species buffers temperature destabilizing effect. Journal of Applied Ecology 59, 2730–2741. doi:10.1111/1365-2664.14267.

Schume, H., Jost, G., Hager, H., 2004. Soil water depletion and recharge patterns in mixed and pure forest stands of European beech and Norway spruce. Journal of Hydrology 289, 258–274. doi:10.1016/j.jhydrol.2003.11.036.

Schwendenmann, L., Pendall, E., Sanchez-Bragado, R., Kunert, N., Hölscher, D., 2015. Tree water uptake in a tropical plantation varying in tree diversity: interspecific differences, seasonal shifts and complementarity. Ecohydrology 8, 1–12. doi:10.1002/eco.1479.

Seynave, I., Bailly, A., Balandier, P., Bontemps, J.D., Cailly, P., Cordonnier, T., Deleuze, C., Dhôte, J.F., Ginisty, C., Lebourgeois, F., Merzeau, D., Paillassa, E., Perret, S., Richter, C., Meredieu, C., 2018. GIS Coop: networks of silvicultural trials for supporting forest management under changing environment. Annals of Forest Science 75, 1–20. doi:10.1007/s13595-018-0692-z.

Toïgo, M., Vallet, P., Perot, T., Bontemps, J.D., Piedallu, C., Courbaud, B., 2015. Overyielding in mixed forests decreases with site productivity. Journal of Ecology 103, 502–512. doi:10.1111/1365-2745.12353.

Trogisch, S., Liu, X., Rutten, G., Xue, K., Bauhus, J., Brose, U., Bu, W., Cesarz, S., Chesters, D., Connolly, J., Cui, X., Eisenhauer, N., Guo, L., Haider, S., Härdtle, W., Kunz, M., Liu, L., Ma, Z., Neumann, S., Sang, W., Schuldt, A., Tang, Z., van Dam, N.M., von Oheimb, G., Wang, M.Q., Wang, S., Weinhold, A., Wirth, C., Wubet, T., Xu, X., Yang, B., Zhang, N., Zhu, C.D., Ma, K., Wang, Y., Bruelheide, H., 2021. The significance of tree-tree interactions for forest ecosystem functioning. Basic and Applied Ecology 55, 33–52. doi:10.1016/j.baae.2021.02.003.

Trumbore, S., Brando, P., Hartmann, H., 2015. Forest health and global change. Science 349, 814–818. doi:10.1126/science.aac6759.

Vidal, J.P., Martin, E., Franchistéguy, L., Baillon, M., Soubeyroux, J.M., 2010. A 50-year high-resolution atmospheric reanalysis over France with the Safran system. International Journal of Climatology 30, 1627–1644. doi:10.1002/joc.2003.

Vilà, M., Vayreda, J., Comas, L., Ibáñez, J.J., Mata, T., Obón, B., 2007. Species richness and wood production: a positive association in Mediterranean forests. Ecology Letters 10, 241–250. doi:10.1111/j.1461-0248.2007.01016.x.

Zeller, L., Pretzsch, H., 2019. Effect of forest structure on stand productivity in Central European forests depends on developmental stage and tree species diversity. Forest Ecology and Management 434, 193–204. doi:10.1016/j.foreco.2018.12.024.

Zhang, Y., Chen, H.Y.H., Reich, P.B., 2012. Forest productivity increases with evenness, species richness and trait variation: a global meta-analysis. Journal of Ecology 100, 742–749. doi:10.1111/j.1365-2745.2011.01944.x.

